# Decoding the activated stem cell phenotype of the vividly maturing neonatal pituitary

**DOI:** 10.1101/2022.01.18.476723

**Authors:** Emma Laporte, Florian Hermans, Silke De Vriendt, Annelies Vennekens, Diether Lambrechts, Charlotte Nys, Benoit Cox, Hugo Vankelecom

## Abstract

The pituitary represents the endocrine master regulator. In mouse, the gland undergoes vivid maturation immediately after birth. Here, we in detail portrayed the stem cell compartment of neonatal pituitary. Single-cell RNA-sequencing pictured an active gland, revealing proliferative stem as well as hormonal (progenitor) cell populations. The stem cell pool displayed a hybrid epithelial/mesenchymal phenotype, characteristic of development-involved tissue stem cells. Organoid culturing recapitulated the stem cells’ phenotype, interestingly also reproducing their paracrine activity. The pituitary stem cell-activating interleukin-6 (IL-6) advanced organoid growth, although the neonatal stem cell compartment was not visibly affected in IL-6^-/-^ mice, likely due to cytokine family redundancy. Further transcriptomic analysis exposed a pronounced WNT pathway in the neonatal gland, shown to be involved in stem cell activation and to overlap with the (fetal) human pituitary transcriptome. Following local damage, the neonatal gland efficiently regenerates, despite absence of additional stem cell proliferation, or upregulated IL-6 or WNT expression, all in line with the already high stem cell activation status, thereby exposing striking differences with adult pituitary. Together, our study decodes the stem cell compartment of neonatal pituitary, exposing an activated state in the vividly maturing gland. Understanding stem cell activation is key to potential pituitary regenerative prospects.

## Introduction

As central hub of the endocrine system, the pituitary plays a quintessential role in governing the endocrine glands throughout the body, thereby regulating key physiological processes including growth, metabolism, fertility, stress and immunity. To execute this primordial function, the pituitary has to properly develop into a compound gland of multiple endocrine cells, encompassing somatotropes (producing growth hormone (GH)), corticotropes (adrenocorticotropic hormone (ACTH)), lactotropes (prolactin (PRL)), gonadotropes (luteinizing hormone (LH) and/or follicle-stimulating hormone (FSH)) and thyrotropes (thyroid-stimulating hormone (TSH)) (Melmed, 2011; Willems et al., 2014). In the mouse, the pituitary undergoes an intense growth and maturation process during the first postnatal weeks, with number and size of hormone-producing cells substantively expanding following proliferation of committed (progenitor) and endocrine cells (Carbajo-Pérez et al., 1990; Laporte et al., 2021; Sasaki, 1988; Taniguchi et al., 2002; Zhu et al., 2015). Simultaneously, the local stem cell (SOX2^+^) compartment shows signs of activation including elevated abundance and expression of stemness pathways when compared to the adult pituitary stem cells (Gremeaux et al., 2012; Laporte et al., 2021). The stem cells can give rise to new endocrine cells, a property which has been found most prominent (although not extensive) during this neonatal period (Andoniadou et al., 2013; Rizzoti et al., 2013; Zhu et al., 2015). In contrast, stem cells in the mature, adult pituitary are quiescent and do not highly contribute to new endocrine cells during the (slow) homeostatic turnover of the gland (Andoniadou et al., 2013; Laporte et al., 2021; Rizzoti et al., 2013; Vankelecom et al., 2014). However, the adult stem cells become activated following local injury in the gland, showing a proliferative reaction and signs of differentiation toward the ablated cells, coinciding with substantial regeneration (Fu et al., 2012). We recently identified interleukin-6 (IL-6) to be upregulated in the adult pituitary (in particular its stem cells) upon this local damage which activated the stem cells (Vennekens et al., 2021). However, it is not clear what molecular mechanisms underlie the activation status of the stem cells in the neonatal gland.

Here, we set out to decode the stem cells’ phenotype during the neonatal pituitary maturation stage starting from single-cell RNA-sequencing (scRNA-seq) interrogations, which uncovered an activation-designating hybrid epithelial-mesenchymal and WNT profile. We applied organoid and mouse models which supported and expanded bioinformatic findings. Moreover, we found that the neonatal pituitary displayed high regeneration efficiency, more pronounced than in older mice (Fu et al., 2012; Willems et al., 2016), which is in line with its activated (stem cell) nature.

Further decoding the molecular mechanisms underlying stem cell activation may open the door toward regenerative approaches for repairing harmed pituitary tissue, having serious endocrine consequences. In this prospect, our single-cell transcriptome database of neonatal (as well as adult) pituitary provides a highly valuable resource.

## Results

### Single-cell transcriptomics pictures vivid developmental activity in the neonatal maturing pituitary

To obtain a granular view on the cell type and activity landscape in the neonatal maturing pituitary, we performed single-cell RNA-sequencing (scRNA-seq) analysis of the major endocrine anterior pituitary (AP) from neonatal mouse (postnatal day 7, PD7), and contrasted the data with adult AP (Vennekens et al., 2021). After quality control and exclusion of doublets and dead and low-quality cells (Figure 1 - figure supplement 1A; (Vennekens et al., 2021)), the neonatal AP cell transcriptomic data were integrated with our recent scRNA-seq dataset of adult AP (Vennekens et al., 2021) followed by filtering-out of ambient RNA (Figure 1A). Unsupervised clustering and annotation using established lineage markers revealed all main endocrine cell types, as well as distinct clusters of pericytes and of stem, mesenchymal, immune and endothelial cells (Figure 1A and B – figure supplement 1B). Of note, the neonatal data also revealed a small contaminating cluster with posterior lobe (PL) signature (i.e. *Nkx2-1*, *Rax* (Cheung et al., 2018) and pituicyte markers *Scn7a*, *Cldn10* (Q. Chen et al., 2020)), in addition expressing the stem cell markers *Sox2* and *Sox9* (Figure 1A and B – figure supplement 1B), the latter in accordance with a previous report (Cheung et al., 2018). In general, sizable overlap between neonatal and adult cell clusters was observed (Figure 1A; visualization using Uniform Manifold Approximation and Projection (UMAP)). In particular, two stem cell subclusters (SC1 and SC2) were discerned in the neonatal AP as before also identified in the adult gland (Vennekens et al., 2021). These subclusters exhibit a diverging transcriptomic fingerprint, with SC1 displaying more prominent expression of keratins *Krt8* and *Krt18*, and SC2 of *Six1* and *Sox2* (Figure 1 - figure supplement 1C), similar to the adult gland (Vennekens et al., 2021). Notwithstanding considerable overlap, clear differences emerged between neonatal and adult gland. First, the stem cell populations (both SC1 and SC2) are larger in neonatal than adult AP (3- to 4-fold; Figure 1A), corroborated *in situ* by immunofluorescence analysis of the SOX2^+^ cells (Figure 1 - figure supplement 1D). Moreover, higher proliferative activity was observed in this SOX2^+^ cell compartment (as assessed by Ki67 immuno-analysis; Figure 1 - figure supplement 1D), as also found before (Gremeaux et al., 2012). In line, the scRNA-seq interrogation exposed a clear proliferative stem cell cluster (Prolif SC; as identified by co-expression of proliferation genes *Mki67, Pcna, Top2a, Mcm6*, with stem cell markers *Sox2, Krt8*, *Sox9*) in the neonatal AP, not discerned in the adult gland (Figure 1A and B – figure supplement 1B). Gene Ontology (GO) analysis using differentially expressed genes (DEGs) indeed revealed enrichment of cell cycle processes in the Prolif SC *versus* the aggregate SC1 and SC2 clusters (Figure 1 - figure supplement 1E; Supplementary file 1A and 2A).

**Figure 1.**
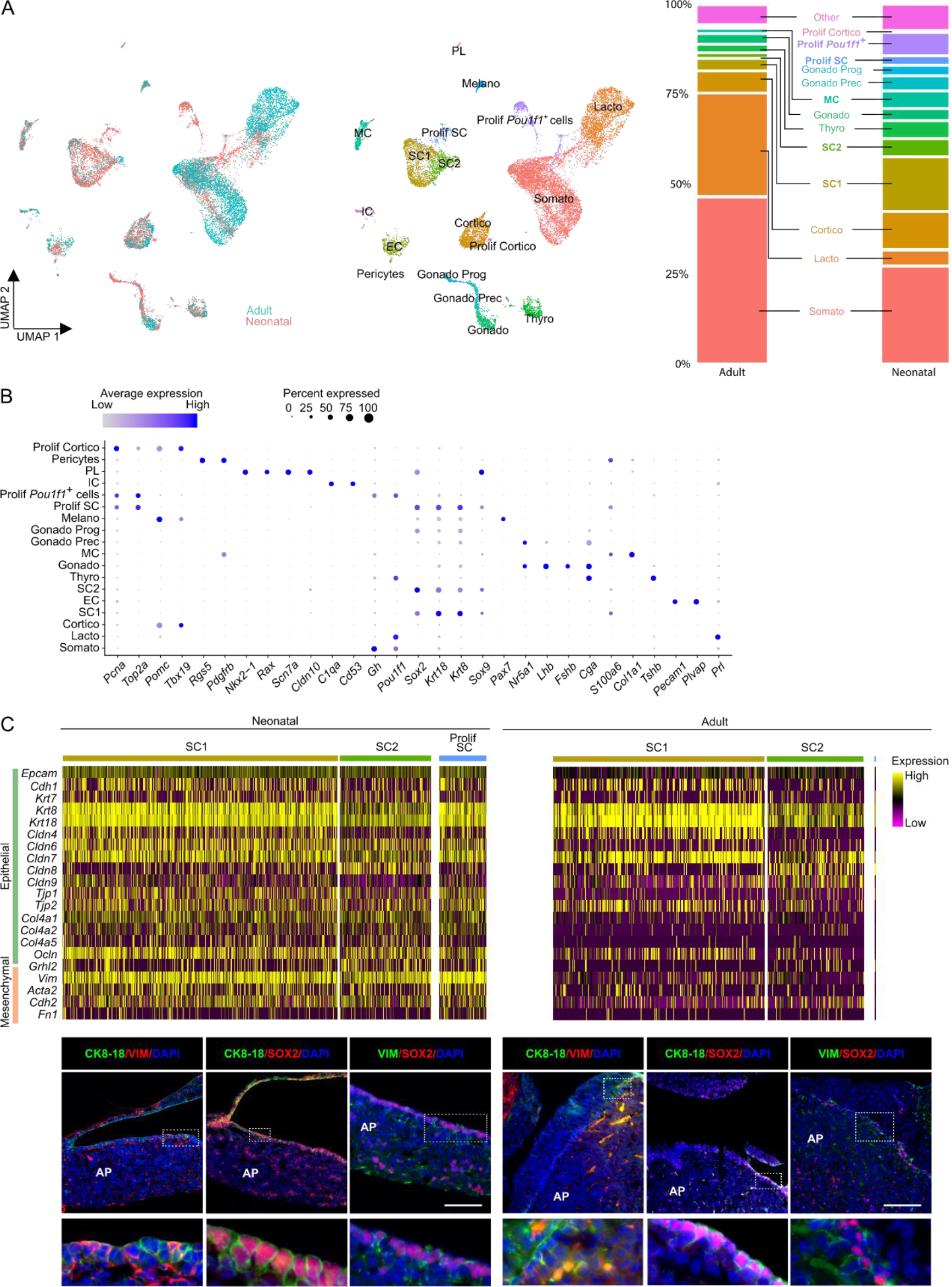
Single-cell transcriptomics of the neonatal maturing pituitary and its activated stem cell compartment. (A) *Left*: UMAP plot of neonatal and adult AP combined. *Middle*: UMAP plot of the annotated cell clusters in the integrated AP samples (i.e. collective single-cell transcriptome datasets from neonatal and adult AP). Somato, somatotropes; Lacto, lactotropes; (Prolif) Cortico, (proliferating) corticotropes; Gonado, gonadotropes; Thyro, thyrotropes; Melano, melanotropes; SC1 and SC2, stem cell cluster 1 and 2; EC, endothelial cells; IC, immune cells; MC, mesenchymal cells; PL, posterior lobe cells; Gonado Prog, gonadotrope progenitor cells; Gonado Prec, gonadotrope precursor cells. *Right*: Bar plots showing proportions of each cell cluster at both ages. (B) Dot plot displaying percentage of cells (dot size) expressing indicated marker genes with average expression levels (color intensity; see scales on top) in the collective AP samples (i.e. adult and neonatal AP). (C) *Top*: Heatmap displaying scaled expression of selected epithelial and mesenchymal marker genes in the stem cell clusters SC1, SC2 and Prolif SC of neonatal and adult AP. *Bottom*: Immunofluorescence staining of CK8/18 (green), VIM (red/green) and SOX2 (red) in neonatal and adult AP. Nuclei are stained with Hoechst33342 (blue). Boxed area is magnified. (Scale bar, 100 µm). The following figure supplement is available for figure 1: Figure supplement 1. Single-cell transcriptomics of the neonatal maturing pituitary and its activated stem cell compartment.

Interestingly, proliferative subclusters were also perceived for other cell types, moreover exclusively in the neonatal gland. First, we discerned a proliferative *Pit1/Pou1f1*^+^ cell group. *Pit1* represents the transcriptional regulator of hormone expression in somatotropes, lactotropes and thyrotropes, and its expression also marks differentiation of progenitors within this so-called *Pit1*^+^ lineage (Zhu et al., 2015). In addition, we distinguished a proliferative corticotrope subcluster. Both findings were corroborated by DEG/GO and hormone/Ki67 immunostaining analyses (Figure 1 - figure supplement 1F; Supplementary file 1B,C and 2B,C). Strikingly, the neonatal AP also contains gonadotrope progenitor and precursor subclusters as characterized by gradually increasing gonadotrope markers (*Nr5a1, Cga, Lhb, Fshb, Gnrhr*) in parallel with declining stem cell factors (*Sox2, Krt8, Sox9*) (Figure 1A and B – figure supplement 1B). The gonadotrope lineage arises last during embryonic development just before birth (Stallings et al., 2016), and some cells in the neonatal AP indeed co-express SOX2 and the common gonadotropin subunit αGSU (Figure 1 - figure supplement 1G). In further support of an endocrine progenitor phenotype (as opposed to a stem cell state), DEG analysis revealed upregulation of genes involved in hormone production (e.g. *Chga, Ascl1, Cga, Insm1*) and enrichment of GO terms related to endocrine differentiation and secretion when compared to the stem cell clusters (Figure 1 - figure supplement 1G; Supplementary file 1D and 2D).

Taken together, scRNA-seq interrogation captured and embodied the vivid developmental activity that is taking place in the maturing neonatal gland, epitomized in the stem cell as well as endocrine lineage phenotypes. While the hormonal cell types (particularly somatotropes and lactotropes) rise in abundance toward adulthood, stem cells as well as supportive mesenchymal cells (MC) are more numerous in the neonatal gland (Figure 1A), suggesting that the latter populations play an active role in the critical development and growth of the gland at this acute period after birth, which becomes less prominent at mature age.

### Neonatal pituitary stem cells display a hybrid epithelial/mesenchymal phenotype

Using the scRNA-seq dataset, we assessed the epithelial character of the stem cell clusters. Surprisingly, the stem cell compartment of the neonatal AP displays a highly mixed epithelial/mesenchymal (E/M) phenotype, manifestly expressing both epithelial and mesenchymal markers, which is clearly faded in the adult gland (Figure 1C). In line, cells that co-express the epithelial markers cytokeratin 8 and 18 (CK8/18) and the mesenchymal marker vimentin (VIM) were found in the marginal-zone (MZ) SOX2^+^ stem cell niche of the neonatal pituitary, and were visibly less prominent in the adult gland (Figure 1C). Similarly, co-expression of SOX2 with VIM was more pronounced in the neonatal gland (Figure 1C). In other developing tissues, stem/progenitor cells also show a hybrid E/M character, indicative of their activation and participation in the tissues’ developmental process (Dong et al., 2018). Recently, a hybrid E/M phenotype was also uncovered in the stem/progenitor cell cluster of the fetal human pituitary, with the mesenchymal aspect lowering along further maturation (Zhang et al., 2020). Moreover, epithelial-to-mesenchymal transition (EMT) has been reported to play a role in mouse pituitary development through driving the exit of stem cells from the MZ toward the developing AP (Davis et al., 2016; Yoshida et al., 2016).

### Organoid culturing recapitulates the neonatal pituitary stem cell phenotype

To further explore the stem cells’ nature of neonatal pituitary, we applied our recently established pituitary organoid model (B. Cox et al., 2019; Vennekens et al., 2021). We have shown that organoids developing from adult mouse AP originate from the SOX2^+^ stem cells and reflect their phenotype and activation state (e.g. higher organoid number following damage; (B. Cox et al., 2019; Vennekens et al., 2021)), thus serving as a valuable pituitary stem-cell biology research model and activation readout tool (B. Cox et al., 2019; Vennekens et al., 2021).

Dissociated AP cells from PD7 mice, embedded in Matrigel droplets (Figure 2A) and cultured in medium previously defined to grow organoids from adult AP (B. Cox et al., 2019) (referred to as pituitary organoid medium or PitOM; Supplementary file 3), were found to generate organoid structures, with a trend of higher number (although not significant; *P*=0.18) than from adult AP (Figure 2B). Surprisingly, individual removal of several growth and signaling factors (i.e. fibroblast growth factors (FGF) 2/8/10, nicotinamide, sonic hedgehog (SHH), the transforming growth factor-β (TGFβ) inhibitor A83-01, the bone morphogenetic protein (BMP) inhibitor noggin and insulin-like growth factor 1 (IGF-1)) from the adult AP-geared PitOM resulted in a visible increase in organoid number developing from neonatal AP (Figure 2 - figure supplement 1A), indicating that these PitOM-included factors actually obstruct organoid outgrowth from the neonatal gland. In contrast, cholera toxin (CT), p38 mitogen-activated protein kinase (MAPK) inhibitor (p38i; SB202190), the WNT-signaling amplifier R-spondin 1 (RSPO1) and epithelial growth factor (EGF) were found essential since organoid formation was largely abrogated in the absence of each one of these factors (Figure 2 - figure supplement 1A). Together, this top-down screening by individually omitting PitOM factors led to an optimized medium for neonatal AP organoid development, further referred to as RSpECT (being the acronym of the essential core factors <u>RS</u>PO1, <u>p</u>38i, <u>E</u>GF and <u>CT</u>; Supplementary file 3).

**Figure 2.**
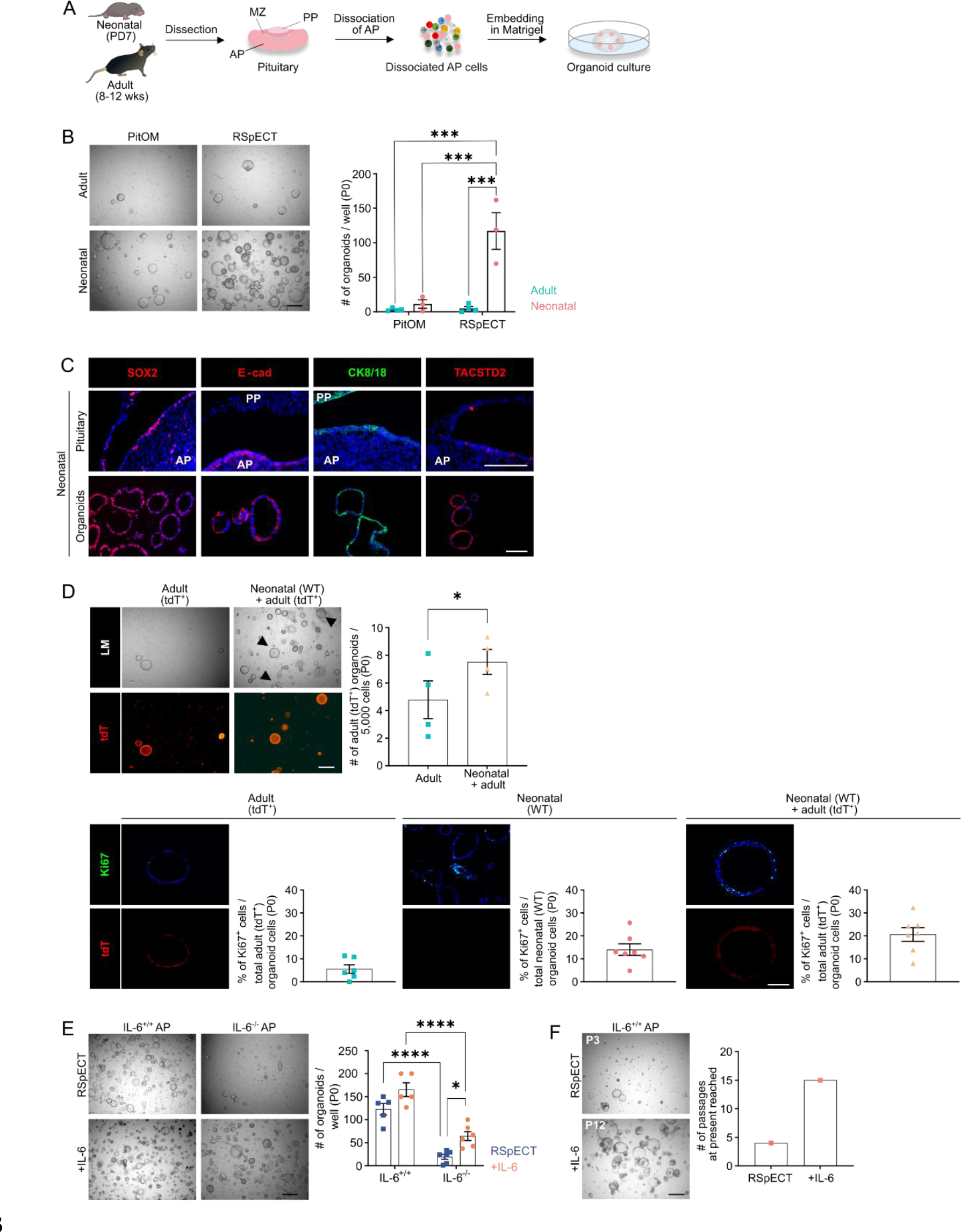
Organoids from neonatal pituitary recapitulate its stem cell phenotype. (A) Experimental schematic for organoid culturing. PD, postnatal day; wks, weeks; MZ, marginal zone; PP, posterior pituitary. Mouse icons obtained from BioRender. (B) Organoid formation efficiency (passage 0, P0) from adult and neonatal AP cultured in PitOM or RSpECT medium. *Left*: Representative brightfield pictures of organoid cultures. (Scale bar, 500 μm). *Right*: Bar graph indicating number of organoids developed per well (mean ± SEM). Data points represent biological replicates. \*\*\**P* ≤ 0.001. (C) Immunofluorescence staining of SOX2, E-cadherin (E-cad), TACSTD2 (all red) and CK8/18 (green) in neonatal pituitary and derived organoids. Nuclei are labeled with Hoechst33342 (blue). (Scale bar, 100 μm). (D) Organoid formation efficiency (P0) starting from AP cells of adult R26^mT/mG^ (tdTomato^+^ (tdT)) mice, or from a 1:1 mixture of adult R26^mT/mG^ and neonatal wild type (WT) AP cells (P0). *Top left*: Light microscopic (LM) and epifluorescence (tdT) pictures. Arrowheads indicate tdT^+^ organoids. (Scale bar, 500 µm). *Top right*: Bar graph indicating number of adult (tdT^+^) organoids developing per 5,000 cells in indicated cultures (mean ± SEM). Data points represent biological replicates. \**P* ≤ 0.05. *Bottom*: Immunofluorescence staining of Ki67 (green) and tdT (red) in AP organoids derived from adult R26^mT/mG^, neonatal WT or a 1:1 mixture of adult R26^mT/mG^ and neonatal WT AP cells. Nuclei are labeled with Hoechst33342 (blue). (Scale bar, 100 μm). Bar graphs showing percentage of Ki67^+^ cells in organoids as indicated (mean ± SEM). Data points represent individual organoids from 3 biological replicates. (E) Organoid formation efficiency from AP of neonatal IL-6^+/+^ and IL-6^-/-^ mice in RSpECT with or without IL-6 (P0). *Left*: Representative brightfield pictures of organoid cultures. (Scale bar, 500 μm). *Right*: Bar plot showing number of organoids developed per well in conditions as indicated (mean ± SEM). Data points represent biological replicates. \**P* ≤ 0.05; \*\*\*\**P* ≤ 0.0001. (F) Neonatal AP organoid passageability with or without IL-6. *Left*: Representative brightfield images at indicated passage. (Scale bar, 500 µm). *Right*: Bars depicting the passage number reached at the end of this study. The following figure supplement is available for figure 2: Figure supplement 1. Organoids from neonatal pituitary recapitulate its stem cell phenotype.

The organoids, as before shown for adult AP (B. Cox et al., 2019), originated from the neonatal tissue-resident SOX2^+^ stem cells. Seeding AP from neonatal SOX2^eGFP/+^ reporter mice (expressing enhanced green fluorescent protein (eGFP) in SOX2^+^ cells) generated only organoids that were fluorescent (eGFP^+^; Figure 2 - figure supplement 1B). Furthermore, when SOX2^eGFP/+^ AP cells were mixed with wildtype (WT) cells, organoids that developed were either eGFP^+^ or not (Figure 2 - figure supplement 1B), thereby pointing to a clonal origin, as further supported by live-culture time-lapse imaging showing organoids growing from individual cells (Figure 2 - figure supplement 1C and Video 1). The obtained organoids display a pituitary stemness phenotype expressing SOX2 as well as other known pituitary stem cell markers (such as E-cadherin, CK8/18, TACSTD2 (alias TROP2); (J. Chen et al., 2009; B. Cox et al., 2019; Fauquier et al., 2008; Vennekens et al., 2021)) (Figure 2C, Video 2 and 3), while being devoid of hormone expression (Figure 2 - Figure supplement 1D). Using the optimized RSpECT medium, the number of organoids that developed from neonatal AP was significantly higher than from adult gland (for which RSpECT did not change the formed organoid number) (Figure 2B). Of note, the much larger number (∼40-fold) can only partially be explained by the higher abundance of SOX2^+^ cells in the neonatal AP (∼3-4-fold; see above). Indeed, normalized to the absolute number of SOX2^+^ cells seeded per well, the proportion of organoid-initiating SOX2^+^ cells is ∼10-fold higher in neonatal than adult AP (Figure 2 - figure supplement 1E). Thus, organoid formation efficiency reflects the higher intrinsic activation (or ‘primed’) status of neonatal pituitary stem cells *per se*.

Very recently, it has been reported that stem cells in early-postnatal (PD14) pituitary can function as autocrine and paracrine signaling center, among others stimulating proliferation within the own stem cell compartment (Russell et al., 2021). Interestingly, when co-cultured, neonatal (PD7) AP organoids (as developed from WT mice) elevate the outgrowth of organoids from adult AP (as established from ubiquitously tdTomato(tdT)-expressing R26^mT/mG^ mice) (Figure 2D), coinciding with increased proliferation of the adult AP organoid-constituting stem cells, showing a Ki67^+^ index reaching the one of neonatal AP-derived organoids (Figure 2D).

Taken together, organoids from neonatal AP reflect the stem cell compartment of the gland at this developmental stage regarding its activated and signaling-center phenotype.

One factor that may be responsible for activating the stem cells in the neonatal pituitary is IL-6 which we recently uncovered as pituitary stem cell-activating factor in adult mouse (Vennekens et al., 2021). Intriguingly, gene expression of *Il6* is low in neonatal AP when compared to adult gland (Figure 2 - figure supplement 1F). Similarly, gene-regulatory network (regulon) activity of the IL-6 signal-mediating JAK/STAT member *Stat3* is low as compared to adult gland (analysis using ‘single-cell regulatory network inference’ (SCENIC); (Aibar et al., 2017)) (Figure 2 - figure supplement 1F). Notwithstanding, IL-6 is still produced, and beneficial, for AP organoid development. Indeed, a significantly smaller number of organoids grew from IL-6 knock-out (IL-6^-/-^) *versus* WT neonatal AP, which was (partially) rescued by adding exogenous IL-6 (Figure 2E). Moreover, although IL-6 addition did not significantly increase the number of organoids formed from WT neonatal AP (most likely because the stem cell activation status is already high) (Figure 2E), it strongly enhanced the passageability of the organoid cultures, from 4-5 passages without IL-6 to at least 15 passages (6 months of expansive culture) with the cytokine (Figure 2F), as also observed before for adult AP (Vennekens et al., 2021). Addition of IL-6 is considered to compensate for the decline in endogenous levels that is observed at passaging (Figure 2 - figure supplement 1G). Of note, the organoids robustly retained their stemness phenotype during the long-term expansion, showing consistent stem cell marker expression (and hormone absence) between early (P2) and late (P8-11) passage (Figure 2 - figure supplement 1H). Intriguingly, the *in vivo* lack of IL-6 does not visibly affect the stem cell compartment and its proliferative phenotype (Figure 2 - figure supplement 1I). One possible explanation is that IL-6’s function is *in vivo* taken over by other cytokines when IL-6 is absent from conception. Expression of the IL-6 family members *Il11* and leukemia-inhibitory factor (*Lif*), and of the related cytokine tumor necrosis factor-α (*Tnf)*, was found slightly enhanced in the neonatal IL6^-/-^ AP (Figure 2 - figure supplement 1J). These cytokines, as well as the related IL-1β, did not significantly increase the number of organoids formed but enhanced organoid expandability (Figure 2 - figure supplement 1K), all similar to IL-6 (Figure 2E and F). The effects of TNFα and IL-1β seem to be mediated by IL-6 since these cytokines did not rescue its absence in organoid culturing from IL-6^-/-^ AP (Figure 2 - figure supplement 1L), and increased the expression of IL-6 (as measured in WT AP organoids) (Figure 2 - figure supplement 1M). In contrast, the IL-6 family members LIF and IL-11, known to act through the same gp130/JAK-STAT pathway (Rose-John, 2018), could substitute for IL-6 since increasing the number of organoids from IL-6^-/-^ AP while not affecting *Il6* expression in WT AP organoids (Figure 2 - figure supplement 1L and M). An important underlying role of JAK-STAT signaling in the organoid development is supported by the strong reduction when adding the STAT3 inhibitor STATTIC (Figure 2 - figure supplement 1N). Taken together, the adult pituitary stem cell activator IL-6, although not highly expressed in neonatal AP stem cells, still advances organoid culturing from neonatal AP but appears not absolutely required for *in situ* stem cell maintenance and proliferative behavior in the neonatal gland, potentially due to redundancy of other IL-6 family cytokines.

### The neonatal pituitary stem cell compartment presents a pronounced WNT pathway

To continue the search for molecular mechanisms underlying the activated stem cell phenotype in the neonatal pituitary, we in further detail compared the neonatal and adult AP scRNA-seq datasets. DEG analysis revealed significantly increased expression of multiple WNT signaling-associated genes (e.g. *Fzd2, Fzd3, Gsk3b, Ctnnb1, Tnks*) in the neonatal *versus* adult stem cell clusters (Figure 3A; Supplementary file 1E). In analogy, GO analysis exposed that biological WNT pathway terms are enriched in the neonatal stem cell compartment (Figure 3B; Supplementary file 2E), and GSEA examination showed significant enrichment of WNT-associated hallmarks (Figure 3 - figure supplement 1A). Regulon activity of the WNT downstream transcription factors *Tcf7l1* and *Tcf7l2* is most prominent in the stem cell clusters (SC1, SC2, Prolif SC) when compared to the endocrine cell clusters (Figure 3C). Besides, high regulon activity is also present in the MC and EC clusters (Figure 3C). Clearly, regulon activity of these WNT transcription factors is higher in neonatal than adult gland (Figure 3C, violin plots). Furthermore, looking at upstream WNT pathway components, we found that ligands and receptors are overall higher expressed at neonatal age as analyzed in the whole AP (Figure 3 - figure supplement 1B). Regarding the stem cell and MC clusters, particularly *Wnt5a* and *Frizzled2* (*Fzd2*) receptor are much higher expressed (both virtually absent in the adult AP) (Figure 3 - figure supplement 1C). This age-related expression difference is even visible in whole AP gene expression analysis (RT-qPCR; Figure 3 - figure supplement 1C). Taken together, the WNT pathway is more prominently present in the neonatal gland, which is proposed to contribute to the activated state of the stem cell compartment. A pronounced WNT profile has recently also been reported in the later PD14 mouse pituitary (Russell et al., 2021).

**Figure 3.**
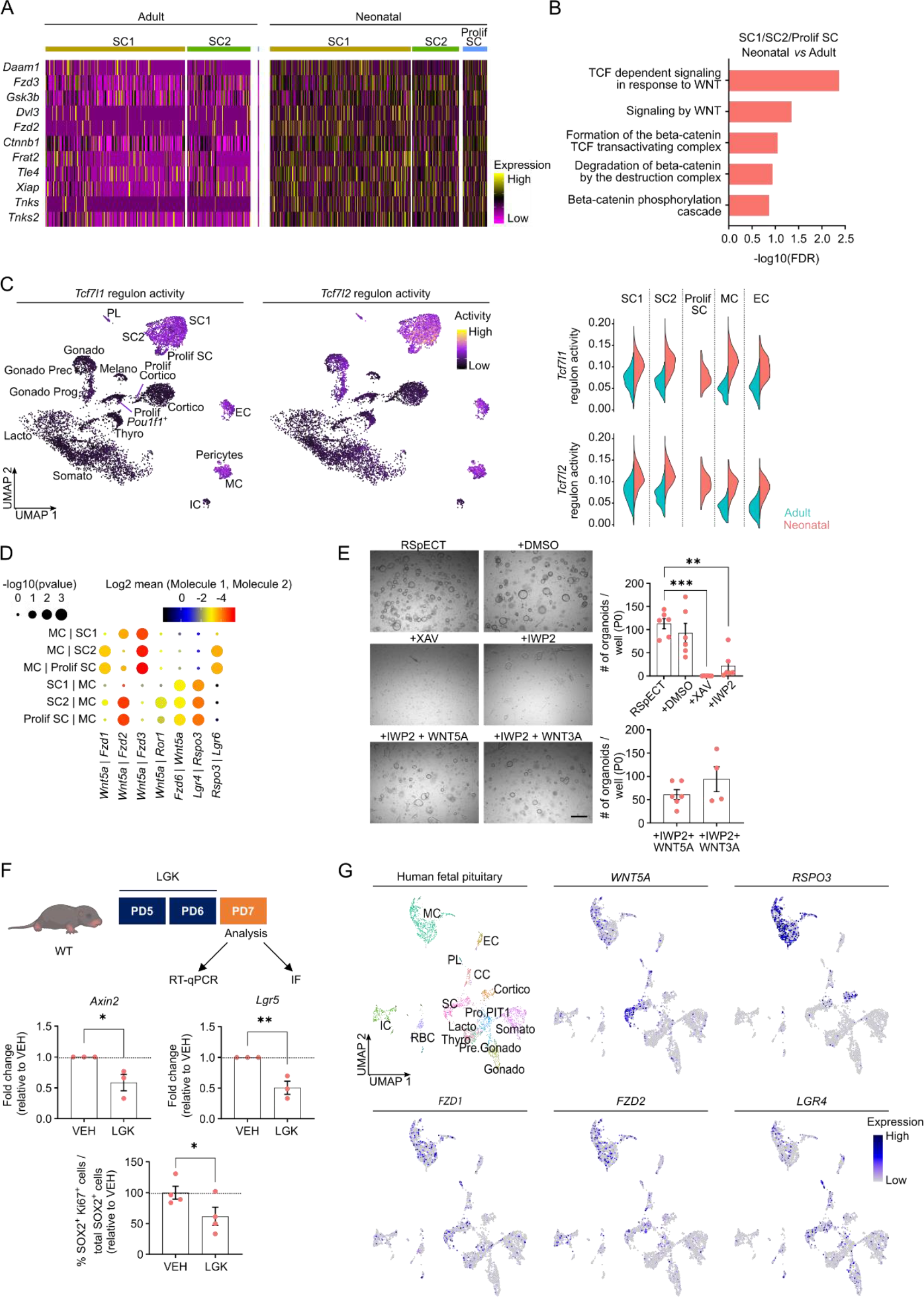
Neonatal pituitary stem cells show a pronounced WNT profile. (A) Heatmap displaying scaled expression of selected WNT-associated DEGs in the stem cell clusters SC1, SC2 and Prolif SC between adult and neonatal AP. (B) DEG-associated GO terms linked with WNT signaling enriched in SC1, SC2 and Prolif SC of neonatal *versus* adult AP. (C) *Left*: *Tcf7l1* and *Tcf7l2* regulon activity projected on UMAP plot of neonatal AP, with indication of cell clusters. *Right*: Violin plots displaying regulon activity of *Tcf7l1* and *Tcf7l2* in the indicated clusters of adult and neonatal AP. (D) Dot plot displaying selected WNT-associated ligand–receptor interactions revealed by CellPhoneDB in neonatal SC1, SC2, Prolif SC and MC clusters. *P* values are indicated by dot size, means of average expression of interacting molecule 1 in cluster 1 and interacting molecule 2 in cluster 2 are specified by color intensity (see scales on top). (E) Organoid development from neonatal AP cells, cultured and exposed to compounds as indicated (P0). *Left*: Representative brightfield pictures of organoid cultures. (Scale bar, 500 μm). *Right*: Bar graphs showing number of organoids formed per well under conditions as indicated (mean ± SEM). Data points represent biological replicates. \*\**P* ≤ 0.01; \*\*\**P* ≤ 0.001. (F) *Top*: Schematic of *in vivo* treatment schedule and analysis. IF, immunofluorescence. Mouse icon obtained from BioRender. *Middle*: Bar plots depicting relative expression level of indicated genes in neonatal AP of mice treated as indicated (relative to vehicle (VEH), set as 1 (dashed line)) (mean ± SEM). *Bottom*: Bar graph showing percentage of SOX2^+^Ki67^+^ cells in SOX2^+^ cell population in neonatal AP of mice treated as indicated (relative to VEH, set as 100% (dashed line)) (mean ± SEM). Data points represent biological replicates. \**P* ≤ 0.05; \*\**P* ≤ 0.01. (G) UMAP plot of the annotated cell clusters in human fetal pituitary (Zhang et al., 2020) and projection of selected WNT-associated genes’ expression. PL, posterior lobe (pituicyte) cells; CC, cell cycle cells; RBC, red blood cells, Pro.PIT1, progenitor cells of PIT1 lineage; Pre.Gonado, precursor cells of gonadotropes. The following figure supplement is available for figure 3: Figure supplement 1. Neonatal pituitary stem cells show a pronounced WNT profile.

We have recently shown that the MC cluster is part of the formerly designated folliculostellate cell population in the (adult) AP (Vennekens et al., 2021), a heterogeneous cell group encompassing, among others, stem cells as well as paracrine-supportive cells (Allaerts et al., 2005; J. Chen et al., 2009; Fauquier et al., 2008; Vennekens et al., 2021). Here, we applied CellPhoneDB (Efremova et al., 2020) on the neonatal scRNA-seq dataset to predict ligand-receptor interactions between MC and stem cells, thereby focusing on the prominently present WNT pathway. Multiple reciprocal interactions of *Wnt5a* with canonical (*Fzd*) and non-canonical (*Ror1*) receptors are projected (Figure 3D), with *Fzd1, Fzd3* and *Fzd6* being the prevalent mediating receptors in the stem cells, and *Fzd2* both in the stem cells and MC (Figure 3D – Figure supplement 1D). Within the RSPO/LGR-driven WNT amplification system, *Rspo3*, being predominantly expressed by MC cells, is forecasted to mainly bind to *Lgr4* and *Lgr6* receptors on the stem cells (Figure 3D - Figure supplement 1D). RNAscope confirmed the pronounced expression of *Lgr4* and *Lgr6* in the *Sox2*^+^ stem cells *in situ*, clearly different from *Rspo3* (Figure 3 - figure supplement 1D).

To functionally validate the proposed stem cell-activating impact of WNT signaling in the neonatal gland, we first applied our *in vitro* organoid model. We observed that the stem-cell WNT pathway components remained expressed in the organoids (Figure 3 - figure supplement 1E). Addition of the WNT inhibitor XAV-939 (XAV; stabilizing the WNT-inhibitory protein AXIN2) at seeding (passage 0 (P0)) completely abolished organoid formation (Figure 3E). Exogenous administration of WNT ligands (WNT5A and WNT3A) to these cultures did not have an effect (Figure 3 - figure supplement 1F), thereby validating the complete blockage of the intracellular (canonical) WNT signaling path by the supplemented XAV. Addition of IWP2 (which blocks endogenous WNT ligand palmitoylation, as needed for secretion from the cell) to the seeded AP cells significantly reduced the number of organoids formed, which was substantially rescued by adding exogenous WNT5A and WNT3A (Figure 3E). Of note, both WNT inhibitors decreased the proliferative activity in the organoids (added to formed organoids at d7 of P0; Figure 3 - figure supplement 1G). Together, these findings show that neonatal AP organoid growth, which reflects stem cell biology/activation, is dependent on (endogenous) WNT activity. Of note, RSPO1, which is typically used in organoid culturing (as we also did here and before, see Supplementary file 3; (B. Cox et al., 2019; Vennekens et al., 2021)), is only slightly expressed in the neonatal pituitary (Figure 3 - figure supplement 1H) whereas *Rspo3* expression is prominent (especially in the MC cluster; Figure 3 - figure supplement 1D). This naturally abundant RSPO ligand was found capable to replace RSPO1 for efficient organoid formation (Figure 3 - figure supplement 1H). Together, our findings advance the concept that the WNT pathway is an activator of pituitary stem cells. In support, addition of WNT5A or WNT3A to cultures from adult AP in which the stem cells are basically not activated (quiescent) (Vennekens et al., 2021) and the WNT pathway is much less prominent (Figure 3A – figure supplement 1A and B), increased stem cell proliferation in the organoids resulting in larger organoid structures (Figure 3 - figure supplement 1I).

To *in vivo* validate our organoid-based finding that the WNT pathway plays a role in stem cell activation in the neonatal pituitary, we treated neonatal (WT) pups with the WNT pathway (porcupine) inhibitor LGK-974 (LGK; Figure 3F). WNT target gene (*Axin2*, *Lgr5*) expression in the AP diminished following LGK administration, thereby verifying its activity and efficacy at the level of the pituitary (Figure 3F). Interestingly, the number of proliferating SOX2^+^ cells decreased (Figure 3F), thus further supporting the involvement of WNT signaling in the activated phenotype of the neonatal AP stem cells.

Finally, as a first translation of our mouse-based pituitary findings to humans, we explored the recently published scRNA-seq dataset of fetal human pituitary (Zhang et al., 2020) regarding WNT component expression. First, we found that the mouse neonatal and fetal human pituitary showed significant overlap and high concordance in clustering outcome (Figure 3 - figure supplement 1J). Of note, *in utero* development of humans is more extended than of mice (Xue et al., 2013), and the neonatal mouse stage thus rather corresponds to (late) embryonic stage in humans. Projection of above specified WNT-associated genes on the human fetal pituitary UMAP plot revealed a comparable expression pattern as in neonatal mouse AP, with, for instance, *WNT5A* being expressed in the SC and MC clusters, and *RSPO3* being most pronounced in the MC cluster (Figure 3G). Thus, the WNT pathway also appears highly present in fetal human pituitary where it may play comparable (e.g. stem-cell activating) roles. Of note, *IL6* is also expressed in the human fetal pituitary stem cells, moreover at similarly low levels as in the neonatal mouse pituitary (Figure 3 - figure supplement 1K).

### The vivid neonatal pituitary shows swift and complete regeneration after local damage

We have previously shown that somatotrope-ablation damage in the adult pituitary triggers acute proliferative activation of the stem cell compartment and expression of GH in the SOX2^+^ cells, and that the somatotrope population was eventually regenerated to 50-60% at 5-6 months after damage infliction (Fu et al., 2012). Here, we investigated whether the neonatal gland, housing a more activated (‘primed’) stem cell compartment, behaves differently regarding acute stem cell reaction and regeneration.

Three-day diphtheria toxin (DT) injection of neonatal (PD4) GH^Cre/+^;R26^iDTR/+^ pups (further referred to as GHCre/iDTR mice, or ‘damaged’ condition) (Figure 4A) resulted in 50-60% ablation of GH^+^ cells (Figure 4B) (as compared to 80-90% ablation at adult age; (Fu et al., 2012)). The stem cell compartment did not visibly react (i.e. no increase in SOX2^+^ cells and their proliferative index, neither proportion-wise nor in absolute cell numbers; Figure 4 – figure supplement 1A), which may be due to their already high activation status. Similarly, organoid formation did not significantly increase (Figure 4 - figure supplement 1B). Fascinatingly, the somatotrope cell population was fully restored to normal numbers, moreover already achieved after 2 months (Figure 4D), meaning a more efficient regenerative capacity of the active neonatal AP than the adult gland (Fu et al., 2012).

**Figure 4.**
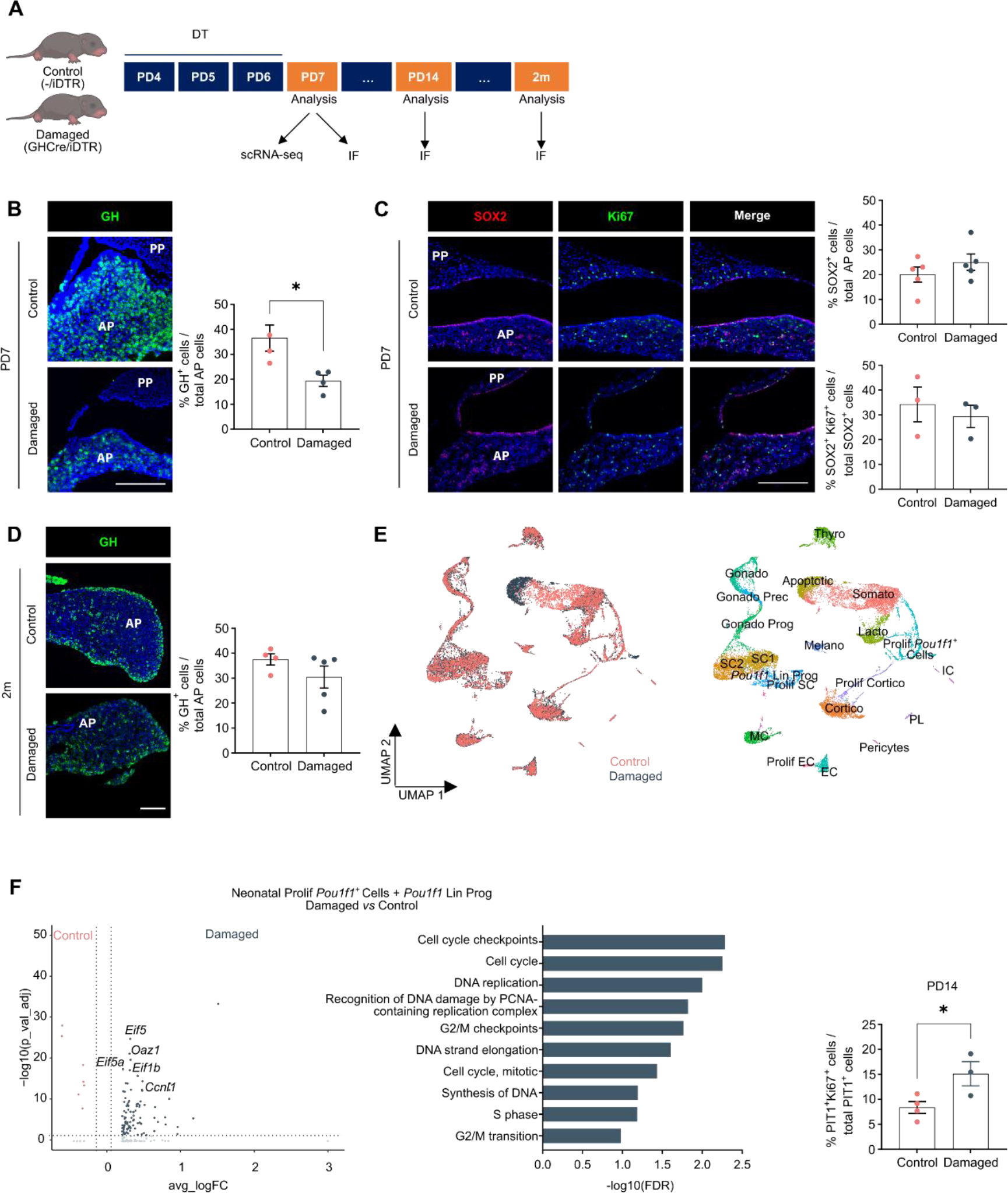
Neonatal pituitary’s reaction to local damage and efficient regeneration. (A) Schematic of *in vivo* treatment schedule and analysis. DT, diphtheria toxin; m, months. Mouse icons obtained from BioRender. (B) Ablation of somatotropes (GH^+^ cells) in neonatal mouse AP. *Left*: Immunofluorescence staining of GH (green) in control and damaged pituitary following DT injection (PD7). Nuclei are labeled with Hoechst33342 (blue). (Scale bar, 100 μm). *Right*: Bar graph showing proportion of GH^+^ cells in AP as indicated (mean ± SEM). Data points represent biological replicates. \**P* ≤ 0.05. (C) SOX2^+^ stem cell reaction to damage. *Left*: Immunofluorescence staining of SOX2 (red) and Ki67 (green) in control and damaged pituitary following DT injection (PD7). Nuclei are labeled with Hoechst33342 (blue). (Scale bar, 100 μm). *Right*: Bar graphs showing proportion of SOX2^+^ cells in AP as indicated, or of SOX2^+^Ki67^+^ cells in SOX2^+^ cell population (mean ± SEM). Data points represent biological replicates. (D) Regeneration of somatotropes in neonatal mouse AP after their ablation. *Left*: Immunofluorescence staining of GH (green) in control and damaged pituitary 2 months after DT-induced ablation. Nuclei are labeled with Hoechst33342 (blue). (Scale bar, 100 μm). *Right*: Bar graph showing proportion of GH^+^ cells in AP as indicated (mean ± SEM). Data points represent biological replicates. (E) *Left*: UMAP plot of control and damaged neonatal AP combined. *Right*: UMAP plot of annotated cell clusters in the integrated neonatal AP samples (i.e. collective single-cell transcriptome datasets from control and damaged AP). (F) *Left:* Volcano plot displaying DEGs in Prolif *Pou1f1^+^* cells and *Pou1f1* Lin Prog clusters from neonatal damaged and control AP. Colored dots represent significantly up- (grey) or down- (orange) regulated genes in damaged *versus* control AP. A selection of cell cycle-associated genes is indicated. *Middle*: DEG-associated GO terms linked with cell cycle processes enriched in Prolif *Pou1f1^+^* cells and *Pou1f1* Lin Prog clusters in neonatal damaged *versus* control AP. *Right*: Bar plot depicting proportion of PIT1^+^ Ki67^+^ cells in PIT1^+^ cell population one week after DT-mediated ablation (PD14) (mean ± SEM). Data points represent biological replicates. \**P* ≤ 0.05. The following figure supplement is available for figure 4: Figure supplement 1. Neonatal pituitary’s reaction to local damage and efficient regeneration.

To start delving into the underlying mechanisms, we performed scRNA-seq analysis of damaged neonatal AP (the day after DT treatment (PD7); Figure 4A), and integrated the data with the control PD7 AP dataset as obtained above (Figure 4E – figure supplement 1C). First, in strong contrast to our recent findings in adult pituitary (Vennekens et al., 2021), *Il6* expression, being very low in neonatal pituitary (see above), was not upregulated in the stem cell and MC clusters following damage (Figure 4 - figure supplement 1D), in line with a lack of extra stem cell activation (while IL-6 activates the stem cells in adult pituitary following damage; (Vennekens et al., 2021)), together indicating that there is a striking difference in adult and neonatal pituitary (stem cell) reaction to damage. Moreover, also in line with the absence of additional stem cell activation, the WNT pathway appeared not to become extra fired upon damage (Figure 4 - figure supplement 1E). Intriguingly, a cluster of *Pou1f1* lineage progenitors (*Pou1f1* Lin Prog) became distinguishable in the aggregate (control + damaged) neonatal AP analysis. DEG and GO examination of the two *Pou1f1* clusters together (Prolif *Pou1f1^+^* cells and *Pou1f1* Lin Prog) revealed enriched cell proliferation genes and terms in damaged *versus* control condition (Figure 4F; Supplementary file 1F and 2F). In line, *in situ* proliferative activity of the PIT1^+^ cells was found increased upon damage (Figure 4F), as well as of already GH-expressing cells (Figure 4 - figure supplement 1F), the latter not observed in adult gland (Fu et al., 2012). Finally, there is also a trend of increase in SOX2^+^GH^+^ cells in damaged *versus* control gland (n=2; Figure 4 - figure supplement 1G), suggesting rising stem cell differentiation. DEG and GO analysis of all SC clusters together revealed enriched NOTCH signaling terms in control *versus* damaged pituitary, or in others words, that NOTCH signaling is downregulated in the stem cells following the inflicted injury (Figure 4 - figure supplement 1H; Supplementary file 1G and 2G), which may be in line with the knowledge that NOTCH inhibition is needed for proper endocrine cell differentiation from stem/progenitor cells as observed during pituitary embryogenesis (B. Cox et al., 2017; Zhu et al., 2006). Taken together, restoration of the GH^+^ cell population in the neonatal pituitary may encompass several mechanisms including proliferative activation of the POU1F1^+^ (progenitor) and derived (or existing) GH^+^ cell populations, and GH/somatotrope differentiation of SOX2^+^ stem cells, together embodying the active nature of the neonatal gland.

In conclusion, these data show that the vividly maturing neonatal gland more efficiently regenerates after injury than the adult gland, but that the already highly activated stem cell population does not show extra visible activation. Further insights into the underlying molecular mechanisms of neonatal AP regeneration will derive from still more extended scRNA-seq and functional explorations.

## Discussion

In our study, we in detail portray the activated nature of the neonatally maturing pituitary, in particular of its stem cells, using single-cell transcriptome mapping and functional *in vitro* (organoid) and *in vivo* (mouse) exploration. The stem cell compartment, shown to be expanded and proliferatively activated in the neonatal gland, presents a pronounced WNT profile. Inhibiting WNT signaling abrogated organoid formation and decreased proliferative activity of the neonatal pituitary stem cell compartment *in vivo*, thereby indicating the importance of the WNT pathway in the activated stem cell phenotype. This WNT signaling may act between the stem cells and supportive mesenchymal cells as projected by CellPhoneDB. Furthermore, the stem cells display a prominent hybrid E/M phenotype in the neonatal gland. A hybrid E/M character has recently also been found in the active stem/progenitor population of the fetal human pituitary (Zhang et al., 2020). More in general, a hybrid E/M nature is present in stem/progenitor cells of developing organs (such as lung, intestine and liver), and underlies their active contribution to the tissues’ development (Dong et al., 2018). Hence, the hybrid E/M phenotype of the neonatal pituitary stem cells likely embodies their active participation in the intense maturation process of the neonatal gland, as proposed during embryogenesis in human pituitary (Zhang et al., 2020). Along this line, we provide further indications that our findings in mouse are translatable to humans. Integrating our neonatal mouse AP scRNA-seq dataset with the recently published data of fetal human pituitary (Zhang et al., 2020) revealed considerable overlap, including an analogous WNT landscape. A pronounced WNT profile has recently also been reported in the early-postnatal (2- weeks-old) mouse pituitary (Russell et al., 2021). Interestingly, this study showed a paracrine- regulatory role of these stem cells, stimulating neighboring committed progenitor cells (as well as stem cells) to proliferate and expand. We also show this paracrine-stimulatory nature of neonatal (1-week-old) pituitary stem cells, inducing proliferative activation of (adult) AP stem cell organoids. Moreover, organoid culturing recapitulated the activated stem cell phenotype of the neonatal AP showing higher outgrowth efficiency. Together, these findings again indicate the faithful reproduction of pituitary stem cell biology and activation by our organoid model.

We recently found that IL-6 is promptly upregulated in the adult pituitary, in particular in its stem cells, following transgenically inflicted damage, associated with proliferative stem cell activation (Vennekens et al., 2021). Intriguingly, IL-6 expression did not rise following damage in the neonatal gland. Together, these findings suggest that the pituitary reacts to damage differently according to developmental age. Along the same line, the dynamic neonatal pituitary more efficiently (faster and more extensively) regenerates than the adult pituitary following the local injury (Fu et al., 2012), likely due to the presence of activated (‘primed’) stem, committed progenitor and just differentiated endocrine cells that all appear to contribute by increased proliferation or differentiation (as supported by transcriptomic and *in situ* immunostaining analyses). Of note, the somewhat lower somatotrope ablation grade in neonatal pituitary may also partly contribute to the higher regeneration level. Interestingly, efficient regenerative capacity is also present in other mouse organs at neonatal age (such as heart and cochlea), while seriously declining or disappearing in adulthood (B. C. Cox et al., 2014; Lam et al., 2018). Detailed unraveling of neonatal-pituitary regenerative mechanisms further needs profound studies, which may find ground in our existing and to be extended single-cell transcriptomic analyses.

IL-6 expression in the neonatal gland is low (much lower than in adult pituitary), and its absence does not visibly affect the neonatally activated stem cell compartment. Either, IL-6 is not needed in the neonatal gland to induce or sustain the activated stem cell phenotype, or its lack has been taken over by cytokine family members such as LIF or IL-11. We provide support for the latter, and also propose that the JAK/STAT pathway may be the common denominator that lies at the basis of stem cell activation in the neonatal gland, hence also critical for organoid outgrowth (as we show with STATTIC and suggest with the IL6^-/-^ mouse) and organoid expandability (as induced by the JAK/STAT-signaling factors IL-6, LIF and IL-11). Adding IL-6 significantly prolonged the passageability of the organoids (as also found before for adult AP-derived organoids; (Vennekens et al., 2021)), thereby compensating for its declining expression at culturing which may be due the stem cells’ removal from the activating (micro-)environment and disappearance of stimulatory factors (such as IL-1β and TNFα which indeed stimulate IL-6 expression). Further and deeper research is now needed to unravel these hypotheses.

In conclusion, our study provides deeper insight into the activated phenotype of the neonatal pituitary stem cell compartment. Together with the scRNA-seq datasets and organoid models developed from adult and aging pituitary (B. Cox et al., 2019; Vennekens et al., 2021), we provide an arsenal of tools to compose a comprehensive view on pituitary stem cell biology, activation and role across key time points of life. Decoding stem cell activation will be needed and instrumental toward future aspirations of repairing and regenerating harmed pituitary tissue.

## Materials and Methods

### Mice and *in vivo* treatments

Mice with C57BL/6 background were used for the experiments, which were approved by the KU Leuven Ethical Committee for Animal Experimentation (P153/2018). Mice were bred and kept in the animal housing facility of the KU Leuven under conditions of constant temperature, humidity and day-night cycle, and had access to food and water *ad libitum*.

Sox2^eGFP/+^ (Sox2^tm1Lpev^) reporter mice contain the gene encoding for eGFP in the *Sox2* open reading frame, resulting in eGFP expression in SOX2-expressing cells (Ellis et al., 2004). Offspring was genotyped for the presence of the *eGFP* transgene by PCR using 5’-TACCCCGACCACATGAAGCA-3’ as forward primer and 5’-TTCAGCTCGATGCGGTTCAC-3’ as reverse primer.

R26^mT/mG^ mice (Gt(ROSA)26Sor^tm4(ACTB-tdTomato,-EGFP)Luo^) contain, in the absence of Cre-mediated recombination, cell membrane-localized fluorescent tdTomato in all cells (Muzumdar et al., 2007). GH^Cre/+^ mice (Tg(Gh1-cre)bKnmn) were crossed with R26^iDTR/iDTR^ animals (Gt(ROSA)26Sor^tm1(HBEGF)Awai^) to create GH^Cre/+^;R26^iDTR/+^ offspring (i.e. heterozygous for both transgenes and abbreviated to GHCre/iDTR) as described in detail before (Fu et al., 2012). Offspring is genotyped for the presence of the *Cre* transgene by PCR using 5’- TGCCACGACCAAGTGACAGCAATG-3’ as forward primer and 5’- ACCAGAGACGGAAATCCATCGCTC-3’ as reverse primer, as previously described (Fu et al., 2012). GHCre/iDTR mice and *Cre*-negative control littermates (further referred to as -/iDTR) (postnatal day (PD) 4) were intraperitoneally (i.p.) injected with 4 ng diphtheria toxin (DT; Merck, Darmstadt, Germany) per g bodyweight, twice a day (8 h in between the injections) for 3 consecutive days. Pituitaries (damaged and control) were isolated and analyzed the following day (PD7), one week later (PD14) or 2 months later.

IL-6^-/-^ (Il6^tm1Kopf^) mice carry a targeted disruption of the *Il6* gene through replacement of the second exon by a neo^r^ cassette (Kopf et al., 1994). These mice were generously provided by Dr. P. Muñoz Cánoves (Cell Biology Unit, Pompeu Fabra University, Barcelona, Spain). Offspring was genotyped for the presence of the neo^r^ cassette and the WT *Il6* gene by PCR using 5’- TTCCATCCAGTTGCCTTCTTGG-3’ as common forward primer, 5’-TTCTCATTTCCACGATTTCCCAG-3’ as WT reverse primer and 5’-CCGGAGAACCTGCGTGCAATCC-3’ as mutant reverse primer.

WT (C57BL/6) neonatal mice (PD5) were treated twice with 5 μg LGK-974 (Biogems, Westlake Village, CA) per g bodyweight or vehicle (corn oil, Merck) trough oral gavage, for 2 consecutive days. Pituitaries were isolated and analyzed the following day (PD7).

### Single-cell RNA-sequencing analysis

The AP of neonatal (PD7) mice was isolated and dispersed into single cells using trypsin (Thermo Fisher Scientific, Waltham, MA), all as previously described (Denef et al., 1978; Van der Schueren et al., 1982). The eventual cell suspension was then subjected to scRNA-seq analyses (2 biological replicates). Cells were loaded on a 10x Genomics cartridge according to the manufacturer’s instructions based on 10x Genomics’ GemCode Technology (10x Genomics, Pleasanton, CA). Barcoded scRNA-seq libraries were prepared with the Chromium Single-cell 3’ v2 Chemistry Library Kit, Gel Bead & Multiplex Kit and Chip Kit (10x Genomics). The libraries were sequenced on an Illumina NextSeq and NovaSeq6000. Data are accessible from ArrayExpress database (accession number E-MTAB-11337). Raw sequencing reads were demultiplexed, mapped to the mouse reference genome (mm10) and gene expression matrices were generated using CellRanger (v3; 10x Genomics). Downstream analysis was performed in R (v.3.6.1) using Seurat (v.3.1.3) (Butler et al., 2018). First, low-quality/dead cells and potential doublets (i.e. with less than 750 genes or more than 8,000 genes and more than 17.5% mitochondrial RNA; see cut offs in Figure 1 - figure supplement 1A) were removed, resulting in a total of 21,419 good-quality single cells (i.e. 9,618 from undamaged AP and 11,801 from damaged AP) for downstream analyses. To allow integrated, comparative examination of undamaged and damaged AP samples, the standard Seurat v3 integration workflow was followed (Stuart et al., 2019). In short, after normalization and identification of variable features for each sample, integration anchors were identified using the FindIntegrationAnchors function with default parameters and dims = 1:20, and data were integrated across all features. Following integration, expression levels were scaled, centered and subjected to principal component analysis (PCA). The top 20 PCs were selected and used for UMAP dimensionality reduction (McInnes et al., 2018). Clusters were identified with the FindClusters function by use of the shared nearest neighbor modularity optimization with a clustering resolution set to 1. This analysis resulted in the identification of 33 distinct clusters, which were annotated based on canonical (AP) cell markers and on previous mouse pituitary scRNA-seq reports (Cheung et al., 2018; Ho et al., 2020; Mayran et al., 2019; Vennekens et al., 2021). Background (ambient) RNA was removed using SoupX (v.1.4.5) with default parameters (Ho et al., 2020). The global contamination (ambient RNA) fraction was estimated at 1.4%, well within the common range of 0-10%. The SoupX-filtered expression matrices were then loaded into Seurat, and processed using standard preprocessing (normalization, variable feature selection and scaling) to enable further downstream analyses.

To integrate our neonatal pituitary dataset with our previously published adult pituitary dataset (Vennekens et al., 2021), we applied Seurat’s reference-based integration approach, for which the ‘adult’ condition was used as reference dataset when applying the FindIntegrationAnchors function (as described above; (Stuart et al., 2019)). The top 30 PCs were selected and used for UMAP dimensionality reduction (McInnes et al., 2018). Clusters were identified with the FindClusters function with a clustering resolution set to 1.6. This analysis resulted in the identification of 37 distinct clusters, which were annotated.

Differential gene expression analysis was performed using the FindMarkers function with default parameters. GO analysis of biological processes was executed on significant differentially expressed genes (DEGs; FDR ≤ 0.05 and logFC ≥ 0.25) using Reactome overrepresentation analysis v3.7 and GOrilla (Eden et al., 2009; Fabregat et al., 2018). Gene-set enrichment analysis (GSEA; v.4.1.0) was performed using normalized expression data (Mootha et al., 2003; Subramanian et al., 2005). Gene sets (hallmarks) tested were obtained from the Molecular Signatures Database (MSigDB; v.7.2) (Liberzon et al., 2015; Subramanian et al., 2005), and converted to mouse gene signatures using MGI batch query (http://www.informatics.jax.org/batch).

Gene regulatory networks (regulons) were determined in our integrated pituitary dataset using pySCENIC, i.e. SCENIC (v.0.9.15; (Aibar et al., 2017)) in Python (v.3.6.9). Raw expression data were normalized by dividing feature counts of each cell by the total counts for that same cell and multiplying by 10,000 followed by log1p transformation. Next, co-expression modules were generated using GRNboost2 algorithm (v.0.1.3) (Moerman et al., 2019). Subsequently, gene regulatory networks were inferred using pySCENIC (with default parameters and mm10 refseqr80 10kb_up_and_down_tss.mc9nr and mm10 refseqr80 500bp_up_and_100bp_down_tss.mc9nr motif collections) resulting in the matrix of AUCell values that represent the activity of each regulon in each cell. The AUCell matrix was imported into Seurat for further downstream analysis, after which it was integrated using the default integration method, as described above. To generate regulon based UMAP plots, the integrated AUCell matrix was scaled, centered and subjected to PCA analysis and the top 20 PCs were selected for UMAP representation.

For integration of our dataset with the recently published human fetal pituitary scRNA-seq dataset (Zhang et al., 2020), the standard Seurat v3 workflow was used as described above. Following integration, expression levels were scaled, centered and subjected to PCA. The top 30 PCs were selected and employed for UMAP dimensionality reduction.

Interactions between pairwise cell clusters were inferred by CellPhoneDB v.2.1.5, which includes a public repository of curated ligands, receptors and their interactions (Efremova et al., 2020). We ran the CellPhoneDB framework using a statistical method and detected ligand-receptor pairs that were expressed in more than 20% of cells. Selected significant ligand-receptor pairs (*P*-value ≤ 0.05 and mean value ≥ 0.5) are shown.

### Organoid culture and treatment

Organoids were developed and cultured as in detail described before (B. Cox et al., 2019). In short, AP cells were plated at a density of 10,000 cells per 30 µl drop of growth factor-reduced Matrigel (Corning, New York, NY) mixed with serum-free defined medium (SFDM; Thermo Fisher Scientific) in a 70:30 ratio (Figure 2A). For all cultures, PitOM or RSpECT (Supplementary file 3) was used unless otherwise stated. At seeding of the primary AP cells and at re-plating of the organoid fragments (i.e. passaging), ROCK inhibitor (Y-27632; 10 µM; Merck) was added to the medium. Organoid cultures were passaged every 10-14 days; the organoids were incubated with TrypLE Express (Thermo Fisher Scientific) and mechanically dispersed until organoid fragments were obtained which were re-seeded in Matrigel drops as above.

To explore their effect on organoid culturing, IL-6 (20 ng/ml; Peprotech, London, UK), IL-1β (10 ng/ml; Peprotech), TNFα (20 ng/ml; Peprotech), LIF (25 ng/ml; Peprotech), IL-11 (25 ng/ml; Peprotech) or STATTIC (20 µM; Merck) was added to the medium.

To assess WNT pathway involvement in organoid culturing, IWP2 (4 µM; Merck), XAV-939 (10 µM; Merck), WNT3A (200 ng/ml; R&D systems, Minneapolis, MN), WNT5A (100 ng/ml; AMSBIO, Cambridge, MA) or RSPO3 (200 ng/ml; Peprotech) were supplemented to the medium.

Brightfield and fluorescence pictures of organoid cultures were recorded using an Axiovert 40 CFL microscope (Zeiss, Oberkochen, Germany). Organoid-forming efficiency was determined by quantifying the number of clearly developed organoids (≥ 100 μm) in whole organoid culture drops using Fiji (https://imagej.net/Fiji; (Schindelin et al., 2012)).

### Histochemical and immunostaining analysis

Pituitary and organoids were fixed in 4% paraformaldehyde (PFA; Merck) and embedded in paraffin using the Excelsior ES Tissue Processor (Thermo Fisher Scientific). Sections were subjected to immunofluorescence staining as described earlier (B. Cox et al., 2019). Antigen retrieval in citrate buffer (Merck) was followed by permeabilization with Triton X-100 (Merck) and blocking with donkey serum (Merck). Following incubation with primary and secondary antibodies (Supplementary file 4), sections were covered with ProLong Gold (Thermo Fisher Scientific) after nuclei counterstaining with Hoechst33342 (Merck).

To quantify SOX2^+^, Ki67^+^, PIT1^+^ and GH^+^ cells, dissociated AP cells were spun onto SuperFrost glass slides (Thermo Fisher Scientific) and the cytospin samples immunostained as described before (Fu et al., 2012). Proportions of immunoreactive cells were counted using Fiji software as previously described (Vennekens et al., 2021).

Images were recorded using a Leica DM5500 upright epifluorescence microscope (Leica Microsystems, Wetzlar, Germany) accessible through the Imaging Core (VIB, KU Leuven) and converted to pictures for figures with Fiji imaging software.

### Gene expression analysis

Total RNA was isolated using the RNeasy Micro kit (Qiagen, Hilden, Germany) and reverse-transcribed (RT) with Superscript III First-Strand Synthesis Supermix (Invitrogen, Waltham, MA). SYBR Green-based quantitative ‘real-time’ PCR (RT-qPCR) was performed, using specific forward and reverse primers (Supplementary file 5), as described before (B. Cox et al., 2019). β-actin (*Actb*), displaying stable expression levels among the conditions tested, was used as housekeeping gene for normalization. Normalized gene expression levels are shown as bar graphs of dCt values (Ct target – Ct housekeeping gene), or gene expression levels were compared between sample and reference as relative expression ratio (fold change) using the formula 2^-(dCt sample - dCt reference)^.

### Time-lapse recording of organoid development and growth

AP cells were seeded at a density of 1,000 cells in 5 µl Matrigel/SFDM drops in 96-well plates (Corning). Brightfield time-lapse images were recorded with the IncuCyte S3 (Sartorius, Göttingen, Germany) every 3 h for 12 days. Time-lapse videos were generated with the IncuCyte software using 10 frames per second.

### 3D imaging of cleared organoids

Whole organoids were immunofluorescently stained and imaged as described (Dekkers et al., 2019). In short, organoids were removed from the Matrigel droplet and fixed in 4% PFA. Permeabilization and blocking were performed with Triton X-100 and bovine serum albumin (BSA; Serva, Heidelberg, Germany) prior to sequential incubation with primary and secondary antibodies (Supplementary file 4). Clearing was achieved by incubating the organoids in a fructose-glycerol solution (Merck; Thermo Fisher Scientific). Samples were mounted and images recorded using a Zeiss LSM 780 – SP Mai Tai HP DS accessible through the Cell and Tissue Imaging Cluster (CIC; KU Leuven). Acquired z-stacks were imported into Fiji image analysis software and 3D reconstructions made using the 3D viewer plugin (Pietzsch et al., 2015).

### RNAscope *in situ* hybridization

Whole pituitary was fixed in 4% PFA for 24 h at room temperature and then paraffin-embedded as described above. Five-µm sections were subjected to *in situ* hybridization with the RNAscope Multiplex Fluorescent Reagent Kit v2 (Advanced Cell Diagnostics, Newark, CA) following the manufacturer’s instructions. Differently labeled RNAscope probes (Advanced Cell Diagnostics) were used for mouse *Sox2* (401041-C3), *Rspo3* (483781-C2), *Lgr4* (318321-C2) and *Lgr6* (404961- C2). Sections were counterstained with DAPI, mounted with ProlongGold, and analyzed with the Zeiss LSM 780 – SP Mai Tai HP DS. Recorded images were converted to pictures using Fiji.

### Electrochemiluminescent measurement of IL-6

IL-6 protein levels were measured with the sensitive electrochemiluminescent ‘Meso Scale Discovery’ (MSD) V-PLEX mouse IL-6 kit (MSD; Rockville, MD), according to the manufacturer’s protocol, in organoid culture supernatant which was collected and centrifuged for 10 min at 1,500 rpm (4°C). Plates were run on the MESO QuickPlex SQ 120 reader and data were analyzed using the MSD discovery workbench software (v4.0.12).

### Statistical analysis

Statistical analysis was performed (when n ≥ 3) using Graphpad Prism (v9.1.2; GraphPad Software, San Diego, CA). (Un-)paired two-tailed Student’s t-test was applied for comparison of 2 groups. In case of multiple comparisons (more than 2 groups), one-way analysis of variance (ANOVA) was done followed by Dunett test when considering 1 independent variable, and two-way ANOVA followed by Sidak’s test in case of 2 independent variables. Statistical significance was defined as *P* < 0.05.

## Acknowledgments

We thank Y. Van Goethem and V. Vanslembrouck for valuable technical help. We are also grateful to the Imaging Core (VIB, KU Leuven) and the CIC (KU Leuven) for use of microscopes and the Center for Brain & Disease Research (CBD) Histology unit (VIB, KU Leuven) for use of histology equipment. We are indebted to Dr. Pura Muñoz-Cánoves (Universitat Pompeu Fabra, Barcelona, Spain) for generously providing the IL-6^-/-^ mice. We thank Thomas Van Brussel, Rogier Schepers and Bram Boeckx (D.L.’s group, KU Leuven) for technical and bioinformatical support in scRNA-seq experiments. The computational resources used for scRNA-seq analysis were provided by the ‘Vlaams Supercomputer Centrum’, managed by the Fund for Scientific Research (FWO) - Flanders. We also thank the FACS core (KU Leuven) for training and use of the MESO QuickPlex SQ 120 reader. Finally, we recognize the Laboratory of Virology and Chemotherapy (Rega Institute; Dr. D. Daelemans) for experimental help and use of the IncuCyte S3.

## Figure supplements

**Figure 1 – figure supplement 1.**
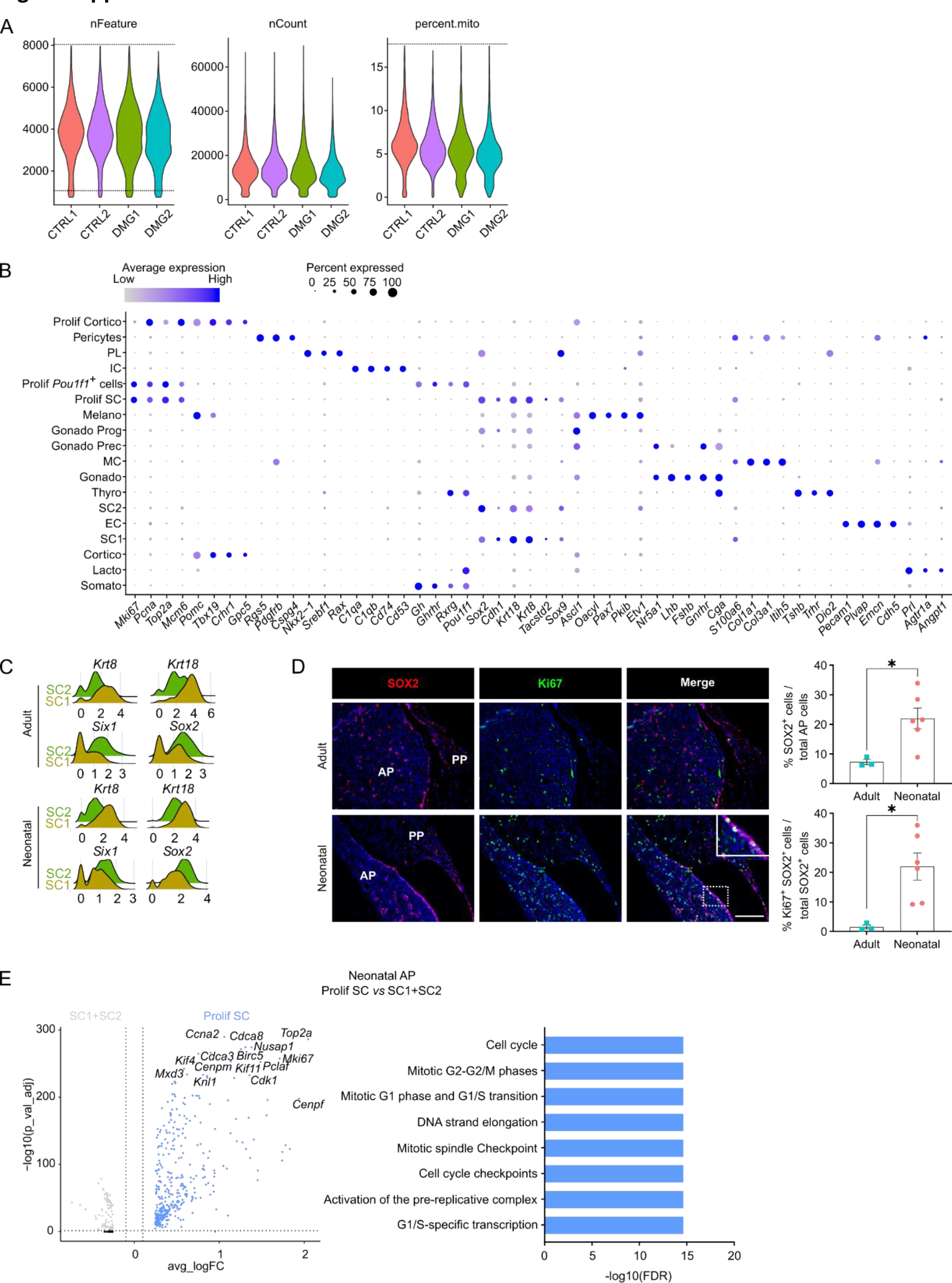
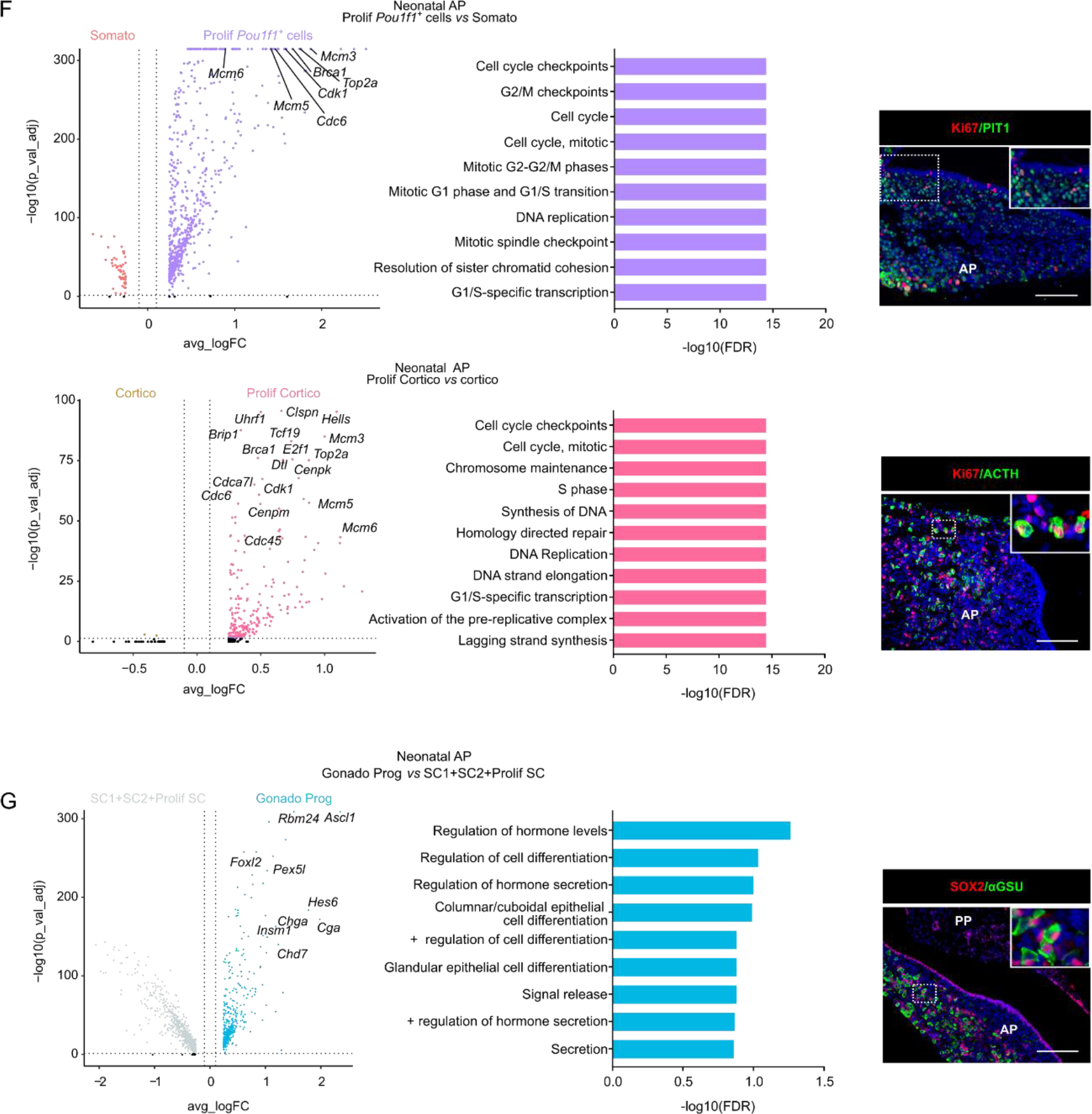
Single-cell transcriptomics of the neonatal maturing pituitary and its activated stem cell compartment. (A) Violin plots showing distribution of number of genes detected per cell (nFeature), total counts per cell (nCount) and percentage of mitochondrial content (percent.mito) per sequenced neonatal AP sample (CTRL, control; DMG, damaged; 2 biological replicates each). Dashed lines indicate cut off values for filtering low-quality/dead cells and doublet exclusion, i.e. removing cells with nFeature below 750 and above 8000 (representing potential doublets) and with percent.mito above 17.5%. (B) Dot plot displaying percentage of cells (dot size) expressing indicated marker genes with average expression levels (color intensity; see scales on top) in the collective AP samples (i.e. adult and neonatal AP). (C) Ridge plot displaying mRNA expression level of indicated genes in SC1 and SC2 of adult and neonatal AP. (D) *Left:* Immunofluorescence staining of SOX2 (red) and Ki67 (green) in adult and neonatal pituitary (separate channels and merge). Nuclei are stained with Hoechst33342 (blue). Boxed area is magnified. (Scale bar, 100 μm). *Right:* Bar graphs showing proportion of SOX2^+^ cells in adult and neonatal AP, or of SOX2^+^Ki67^+^ cells in SOX2^+^ cell population (mean ± SEM). Data points represent biological replicates. \**P* ≤ 0.05. (E) *Left:* Volcano plot displaying DEGs in neonatal AP. Colored dots represent significantly up- (blue) and down- (grey) regulated genes in Prolif SC *versus* SC1 and SC2. A selection of cell cycle-associated genes is indicated. *Right:* DEG-associated GO terms linked with cell cycle processes enriched in Prolif SC *versus* SC1+SC2 of neonatal AP. (F) *Left:* Volcano plot displaying DEGs in neonatal AP. Colored dots represent significantly up- (purple, pink) and down- (orange, yellow) regulated genes in Prolif *Pou1f1^+^* cells *versus* Somato and Prolif Cortico *versus* Cortico, respectively. A selection of cell cycle-associated genes is indicated. *Middle:* DEG-associated GO terms linked with cell cycle processes enriched in Prolif *Pou1f1^+^* cells *versus* Somato or Prolif Cortico *versus* Cortico, respectively, of neonatal AP. *Right:* Immunofluorescence staining of Ki67 (red) and PIT1 or ACTH (both green) in neonatal pituitary. Nuclei are labeled with Hoechst33342 (blue). Boxed areas are magnified. (Scale bar, 100 μm). (G) *Left:* Volcano plot displaying DEGs in neonatal AP. Colored dots represent significantly up- (blue) and down- (grey) regulated genes in Gonado Prog *versus* SC1, SC2 and Prolif SC. A selection of endocrine/secretory-associated genes is indicated. *Middle:* DEG-associated GO terms linked with endocrine/secretory processes enriched in the Gonado Prog *versus* SC1+SC2+Prolif SC of neonatal AP. *Right:* Immunofluorescence staining of SOX2 (red) and αGSU (green) in neonatal pituitary. Nuclei are labeled with Hoechst33342 (blue). Boxed area is magnified. (Scale bar, 100 μm).

**Figure 2 - figure supplement 1.**
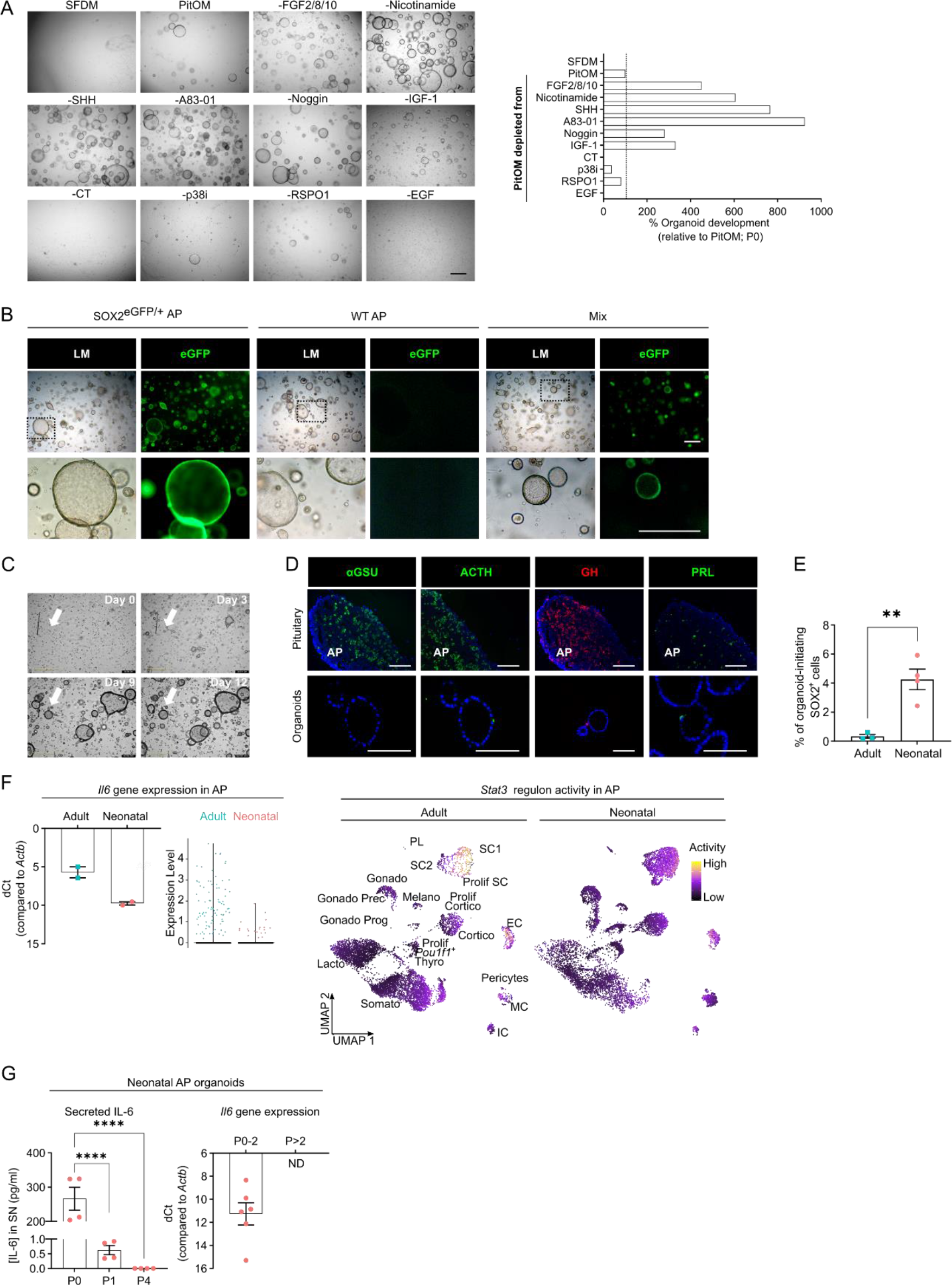
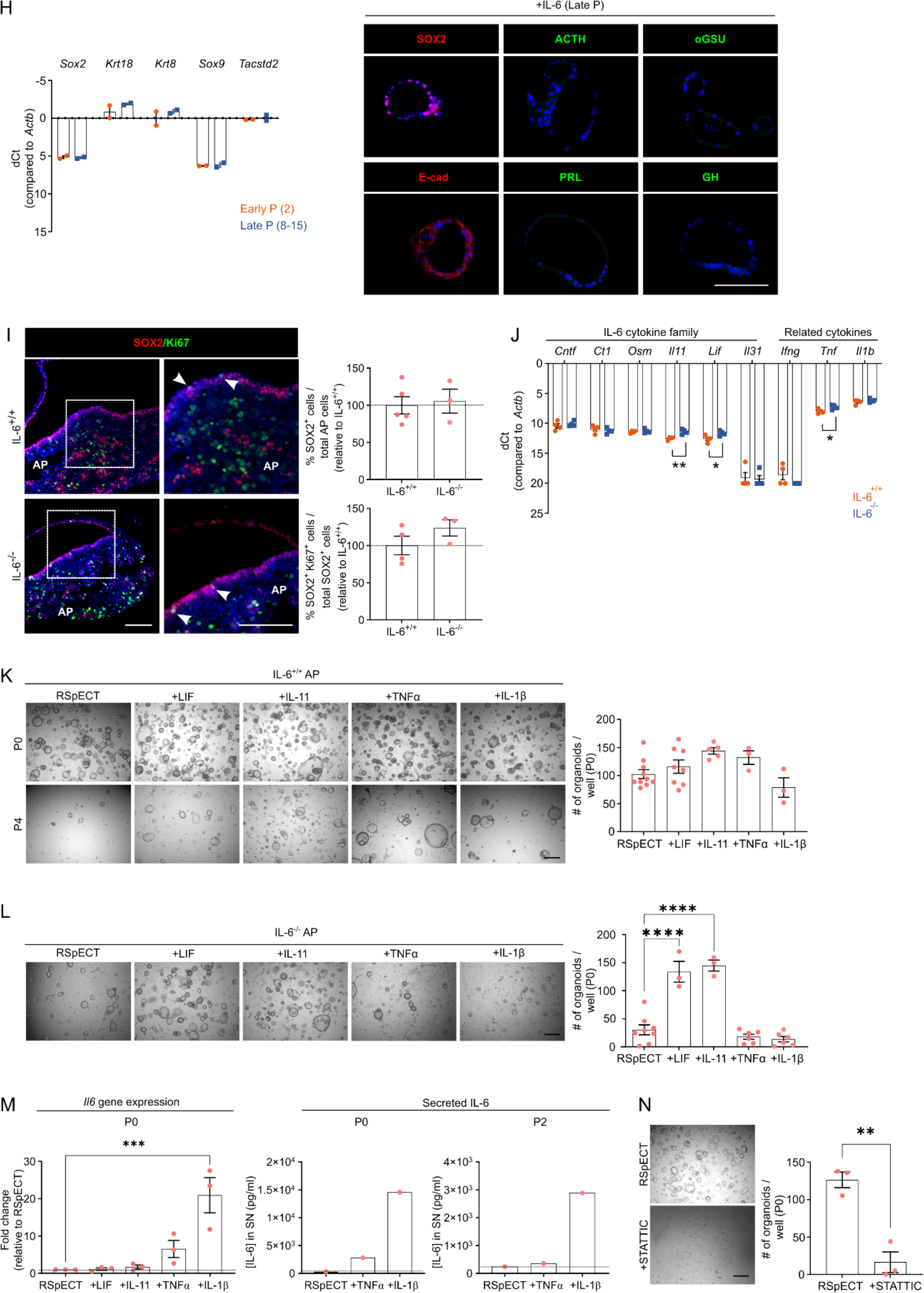
Organoids from neonatal pituitary recapitulate its stem cell **phenotype.** (A) Organoid development from neonatal AP cells in PitOM lacking the indicated factors (P0). *Left:* Representative brightfield pictures of organoid cultures. (Scale bar, 500 μm). *Right:* Bar plot showing percentage organoids developed per well in indicated conditions (relative to PitOM, set as 100% (dashed line)). (B) Organoid formation from AP cells of neonatal SOX2^eGFP/+^ or wildtype (WT) mice, or from a 1:1 mixture (Mix) of neonatal SOX2^eGFP/+^ and WT AP cells (P0). Light microscopic (LM) and epifluorescence (enhanced green fluorescent protein (eGFP)) images are shown. Boxed areas are magnified in the bottom row. (Scale bars, 500 µm). (C) Representative, still brightfield images of live time-lapse recordings (Video 1) of neonatal AP organoid formation (P0) using IncuCyte S3 at days indicated. Arrow points to a single structure at consecutive days of culture, starting from an individual cell. (D) Immunofluorescence staining of αGSU, ACTH, PRL (all green) and GH (red) in neonatal pituitary and derived organoids. Nuclei are labeled with Hoechst33342 (blue). (Scale bar, 100 μm). (E) Bar graph showing percentage of organoid-initiating SOX2^+^ cells per well of 10,000 seeded adult or neonatal AP cells (mean ± SEM). Data points represent biological replicates. \*\**P* ≤ 0.01. (F) *Left:* Bar plot depicting relative *Il6* gene expression level as determined by RT-qPCR (mean ± SEM), and violin plot displaying mRNA expression level of *Il6* in adult and neonatal AP as exposed by scRNA-seq analysis. *Right: Stat3* regulon activity projected on UMAP plot of adult and neonatal AP, with indication of cell clusters. (G) *Left:* Bar graph displaying IL-6 protein levels in supernatant (SN) medium from neonatal AP organoid cultures at indicated passages (mean ± SEM). *Right:* Bar plot depicting relative gene expression level of *Il6* in neonatal AP organoids at indicated passages (mean ± SEM). Data points represent biological replicates. ND, not detectable. \*\*\*\**P* ≤ 0.0001. (H) *Left:* Bar graph showing relative gene expression levels of indicated genes in early (P2) and late (P8-15) passage organoids from neonatal AP (mean ± SEM). Data points represent biological replicates. *Right*: Immunofluorescence staining of SOX2 and E-cad (red) and ACTH, αGSU, PRL and GH (all green) in IL-6-expanded, late passage organoids from neonatal AP. Nuclei are labeled with Hoechst33342 (blue). (Scale bar, 100 μm). (I) *Left:* Immunofluorescence staining of SOX2 (red) and Ki67 (green) in IL-6^+/+^ and IL-6^-/-^ neonatal pituitary. Nuclei are labeled with Hoechst33342 (blue). Boxed areas are magnified. Arrowheads indicate SOX2^+^Ki67^+^ cells. (Scale bar, 100 μm). *Right:* Bar graphs showing proportion of SOX2^+^ cells in IL-6^-/-^ neonatal AP and of SOX2^+^Ki67^+^ cells in SOX2^+^ cell population (relative to IL-6^+/+^ AP, set as 100% (dashed line)) (mean ± SEM). Data points represent biological replicates. (J) Bar graph showing relative gene expression levels of indicated genes in IL-6^+/+^ and IL-6^-/-^ neonatal AP (mean ± SEM). Data points represent biological replicates. \**P* ≤ 0.05; \*\**P* ≤ 0.01. (K) Organoid development from WT (IL-6^+/+^) neonatal AP cells, cultured and exposed to cytokines as indicated (P0). *Left:* Representative brightfield pictures of organoid cultures. (Scale bar, 500 μm). *Right*: Bar graphs showing number of organoids developed per well under the conditions as indicated (mean ± SEM). Data points represent biological replicates. (L) Organoid development from IL-6^-/-^ neonatal AP cells, cultured and exposed to cytokines as indicated (P0). *Left:* Representative brightfield pictures of organoid cultures. (Scale bar, 500 μm). *Right*: Bar graphs showing number of organoids formed per well under the conditions as indicated (mean ± SEM). Data points represent biological replicates. \*\*\*\**P* ≤ 0.0001. (M) *Left:* Bar graph depicting relative gene expression level of *Il6* (relative to expression in RSpECT, set as 1 (dashed line)) (mean ± SEM). Data points represent biological replicates. \*\*\**P* ≤ 0.001. *Right:* IL-6 protein level in SN medium (collected 3 days after seeding/passaging) of neonatal AP organoids in indicated passages and culture conditions (mean ± SEM). Data point represents mean of 2 technical replicates (n = 1). (N) Organoid formation efficiency from neonatal AP in RSpECT with or without STATTIC (P0). *Left:* Representative brightfield pictures of organoid cultures. (Scale bar, 500 μm). *Right*: Bar graph indicating number of organoids developed per well (mean ± SEM). Data points represent biological replicates. \*\**P* ≤ 0.01.

**Figure 3 – figure supplement 1.**
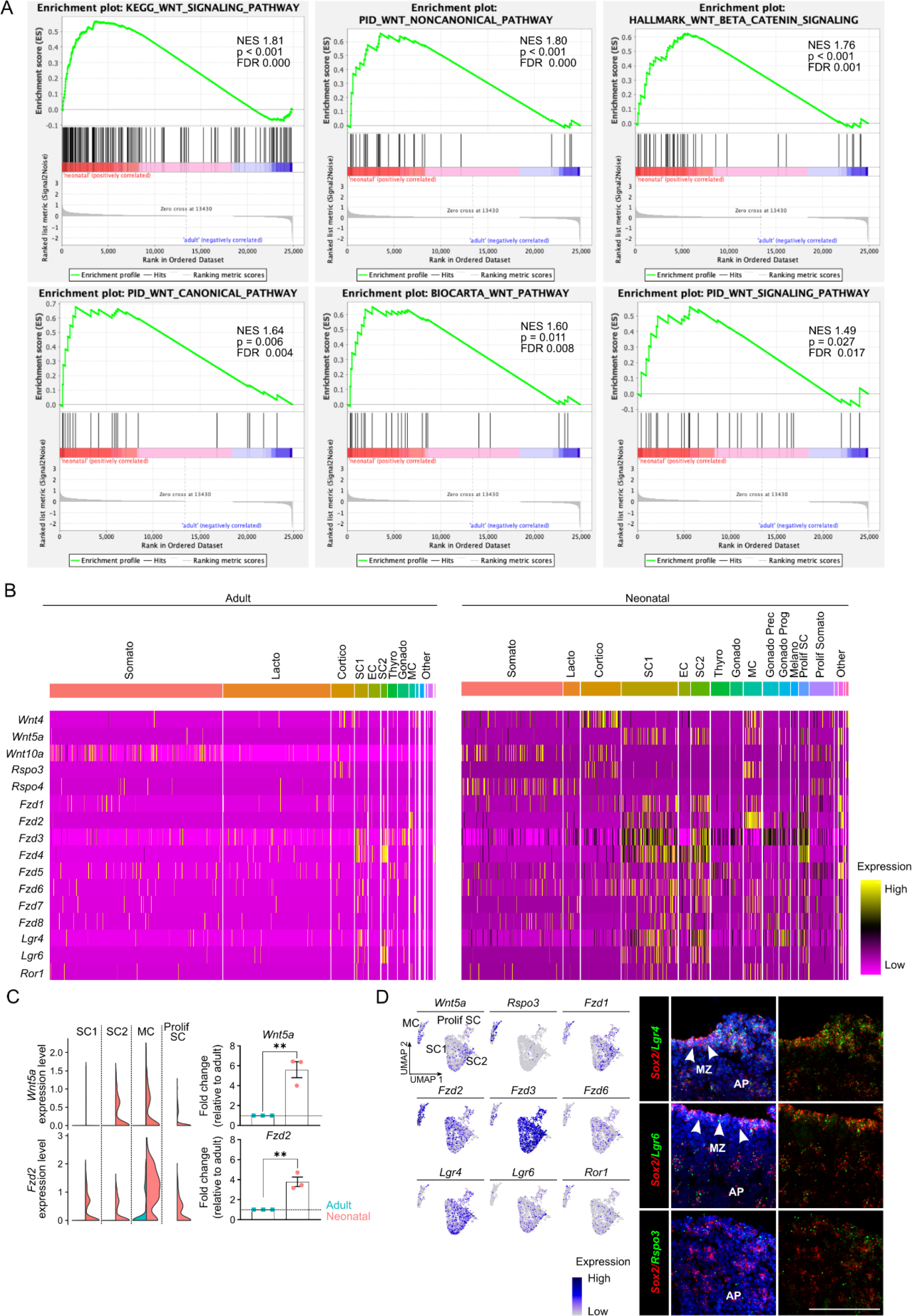
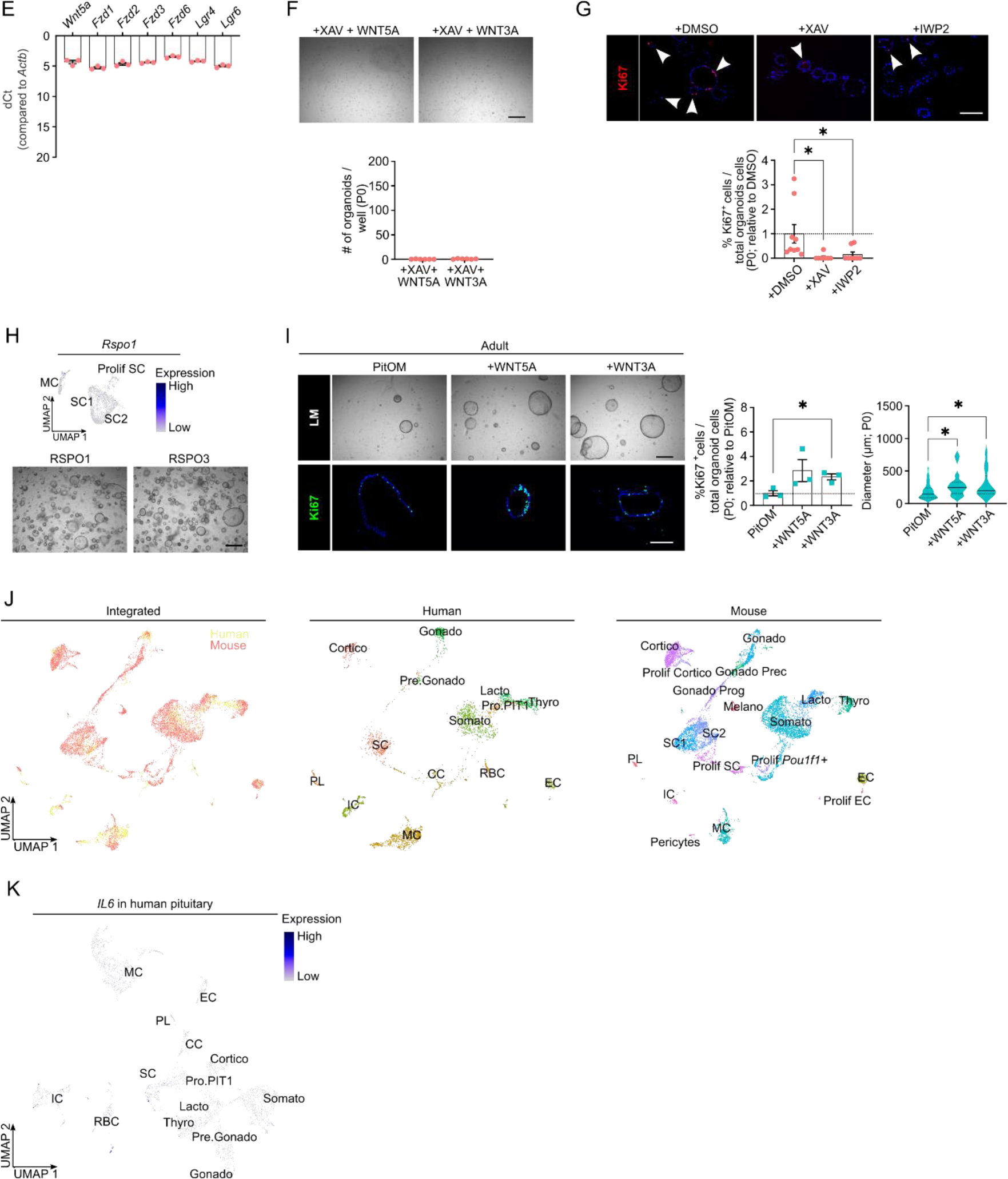
Neonatal pituitary stem cells show a pronounced WNT profile. (A) DEG-based GSEA plots of indicated WNT-related hallmarks in neonatal *versus* adult stem cell clusters (SC1, SC2, Prolif SC). Normalized enrichment score (NES), and *P*- and FDR-values are listed. (B) Heatmaps displaying scaled expression of several WNT ligand and receptor genes in adult and neonatal AP. (C) *Left:* Violin plots displaying mRNA expression level of indicated genes in SC1, SC2, MC and Prolif SC of adult and neonatal AP. *Right:* Bar graphs displaying relative gene expression of indicated genes in neonatal AP, as determined by RT-qPCR (relative to expression in adult AP, set as 1 (dashed line)) (mean ± SEM). Data points represent biological replicates. \*\**P* ≤ 0.01. (D) *Left:* Projection on UMAP plot of selected WNT-associated genes’ expression in neonatal stem cell (SC1, SC2, Prolif SC) and MC clusters, with indication of cell clusters. *Right:* RNAscope *in situ* hybridization analysis of neonatal pituitary for *Sox2* (red), *Lgr4, Lgr6* and *Rspo3* (all green). Nuclei are stained with DAPI (blue). Arrowheads indicate the marginal zone (MZ). (Scale bar, 100 μm). (E) Bar graph displaying relative gene expression level of indicated genes in neonatal AP-derived organoids (mean ± SEM). Data points represent biological replicates. (F) Organoid development from neonatal AP cells, cultured and exposed to WNT ligands as indicated (P0). *Top:* Representative brightfield pictures of organoid cultures. (Scale bar, 500 μm). *Bottom:* Bar graph showing number of organoids formed under conditions as indicated (mean ± SEM). Data points represent biological replicates. (G) *Top:* Immunofluorescence staining of Ki67 (red) in neonatal AP organoids formed under conditions as indicated (P0). Nuclei are stained with Hoechst33342 (blue). Arrowheads indicate Ki67^+^ cells. (Scale bar, 100 µm). *Bottom:* Bar graphs showing percentage of Ki67^+^ cells in organoids as indicated (relative to DMSO, set as 1 (dashed line)) (mean ± SEM). Data points represent individual organoids from 3 biological replicates. \**P* ≤ 0.05. (H) *Top:* Projection on UMAP plot of *Rspo1* gene expression in neonatal stem cell (SC1, SC2, Prolif SC) and MC clusters, with indication of cell clusters. *Bottom:* Organoid development from neonatal AP cells formed in standard RSpECT medium (with RSPO1) or RSpECT medium in which RSPO1 was replaced with RSPO3 (P0). Representative brightfield images are shown. (Scale bar, 500 μm). (I) Organoid development from adult AP cells formed under conditions as indicated (P0). *Top left:* Representative brightfield pictures of organoid cultures. (Scale bar, 500 μm). *Bottom left:* Immunofluorescence staining of Ki67 (green) in adult AP organoids, formed under conditions as indicated. Nuclei are stained with Hoechst33342 (blue). (Scale bar, 100 µm). *Middle:* Bar graph showing percentage of Ki67^+^ cells in organoids as indicated (relative to PitOM, set as 1 (dashed line)) (mean ± SEM). *Right:* Violin plot showing diameter of organoids developed in conditions as indicated. Data points represent biological replicates. \**P* ≤ 0.05. (J) *Left:* UMAP plot of fetal human and neonatal mouse AP combined. *Middle and right:* UMAP plot of annotated cell clusters in human and mouse AP, respectively. Somato, somatotropes; Lacto, lactotropes; Cortico, corticotropes; Gonado, gonadotropes; Thyro, thyrotropes; Melano, melanotropes; SC1 and SC2, stem cell cluster 1 and 2; EC, endothelial cells; IC, immune cells; CT, connective tissue cells; PL, posterior lobe (pituicyte) cells; Gonado Prog, gonado progenitor cells; Gonado Prec, gonadotrope precursor cells; CC, cell cycle cells; RBC, red blood cells, Pro.PIT1, progenitor cells of PIT1 lineage; Pre.Gonado, precursor cells of gonadotropes. (K) Projection of *Il6* gene expression on human fetal pituitary UMAP plot, with indication of cell clusters.

**Figure 4 - figure supplement 1.**
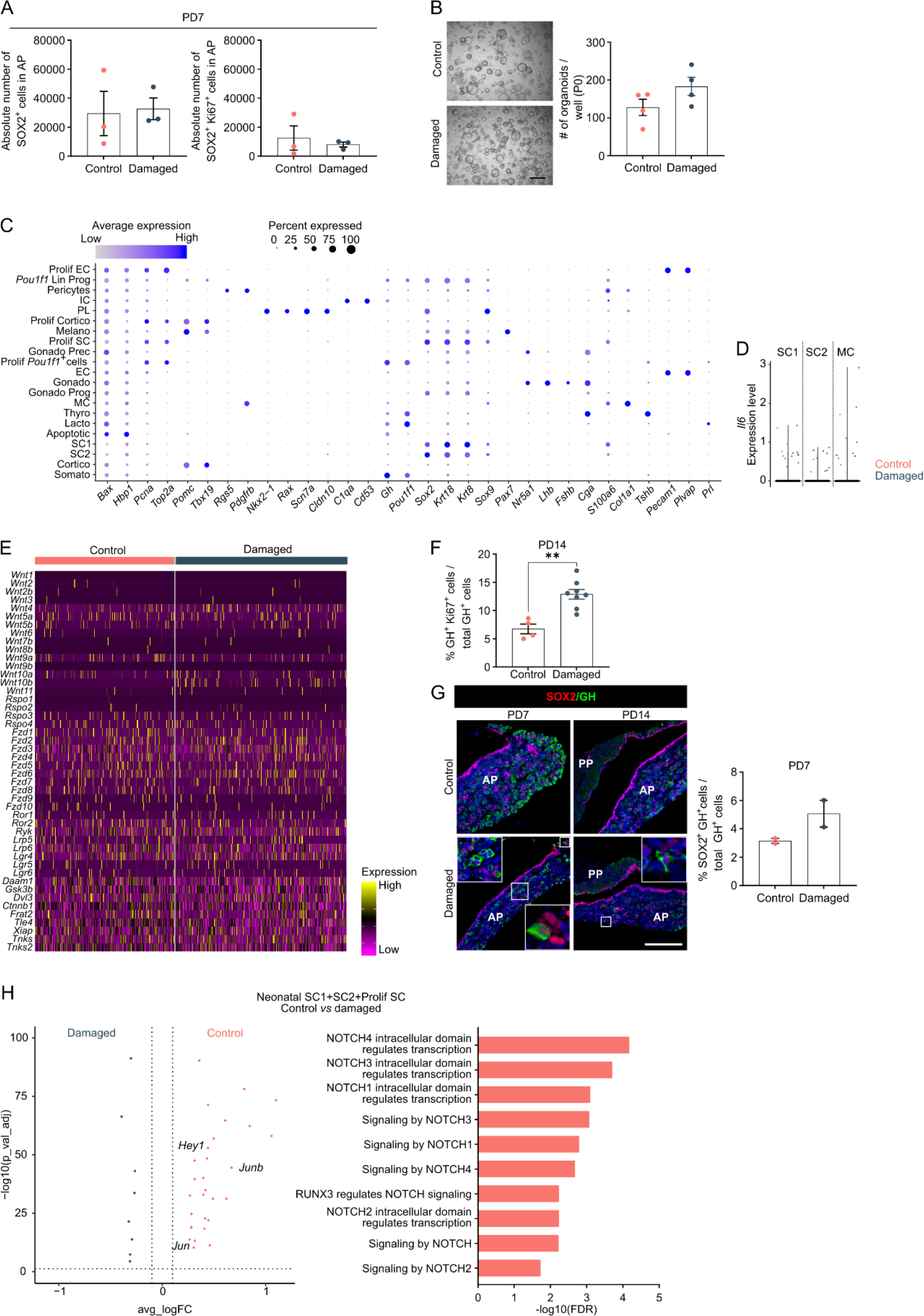
Neonatal pituitary’s reaction to local damage and efficient regeneration. (A) Bar graphs showing absolute number of SOX2^+^ cells in AP as indicated, or of SOX2^+^Ki67^+^ cells in SOX2^+^ cell population following DT injection (PD7) (mean ± SEM). Data points represent biological replicates. (B) Organoid development from control and damaged neonatal AP (P0). *Left:* Representative brightfield pictures of organoid cultures. (Scale bar, 500 μm). *Right:* Bar graph showing number of organoids formed per well (mean ± SEM). Data points represent biological replicates. (C) Dot plot displaying percentage of cells (dot size) expressing indicated marker genes with average expression levels (color intensity; see scales on top) in the collective neonatal AP samples (i.e. control and damaged neonatal AP). (D) Violin plots displaying mRNA expression level of *Il6* in SC1, SC2 and MC of control and damaged neonatal AP. (E) Heatmap displaying scaled expression of selected WNT-associated genes in control and damaged neonatal AP. (F) Bar plot depicting proportion of GH^+^ Ki67^+^ cells within GH^+^ cell population one week after DT-mediated ablation (PD14) (mean ± SEM). Data points represent biological replicates. \*\**P* ≤ 0.01. (G) *Left:* Immunofluorescence staining of SOX2 (red) and GH (green) in control and damaged neonatal pituitary at indicated timepoints after DT-mediated ablation. Nuclei are stained with Hoechst33342 (blue). Boxed areas are magnified. (Scale bar, 100 µm). *Right:* Bar graph showing proportion of SOX2^+^GH^+^ cells in GH^+^ cell population following DT injection (PD7) (mean ± SEM). Data points represent biological replicates. (H) *Left:* Volcano plot displaying DEGs in SC1, SC2 and Prolif SC clusters from neonatal damaged and control AP. Colored dots represent significantly up- (orange) and down- (grey) regulated genes in control *versus* damaged AP. A selection of NOTCH-associated genes is indicated. *Right:* DEG-associated GO terms linked with NOTCH signaling enriched in SC1, SC2 and Prolif SC of neonatal control *versus* damaged AP.

## Legends to supporting files

Video 1

Time-lapse video of neonatal AP organoid formation. Video reconstruction of live time-lapse brightfield images captured by IncuCyte S3 (following the timepoints shown in the timetable on the video) of organoid culture after neonatal AP cell seeding (P0). Cultures were scanned automatically every 3 h for 12 days.

Video 2

3D imaging of neonatal AP organoids. Movie of z-stack through a neonatal AP organoid immunofluorescently stained for SOX2 (red) and E-cadherin (green). Hoechst33342 was used as nuclear stain (blue).

Video 3

3D imaging of neonatal AP organoids. 3D reconstruction of a neonatal AP organoid immunofluorescently stained for SOX2 (red) and E-cadherin (green). Hoechst33342 was used as nuclear stain (blue).

Supplementary file 1

Differentially expressed gene (DEG) analysis. (A) DEG analysis in Prolif SC *versus* SC1+SC2 of neonatal AP. (B) DEG analysis in Prolif *Pou1f1*^+^ cells *versus* Somato of neonatal AP. (C) DEG analysis in Prolif Cortico *versus* Cortico of neonatal AP. (D) DEG analysis in Gonado Prog *versus* SC1+SC2+Prolif SC of neonatal AP. (E) DEG analysis in SC1+SC2+Prolif SC of neonatal *versus* adult AP. (F) DEG analysis in Prolif *Pou1f1*^+^ cells + *Pou1f1* Lin Prog of damaged *versus* control neonatal AP. (G) DEG analysis in SC1+SC2+Prolif SC of control *versus* damaged neonatal AP. The columns represent: gene names (Gene), *P* values (p_val), average log fold change expression (avg_logFC; positive values indicate upregulation and negative values downregulation in first mentioned cluster), percentage of cells expressing the indicated gene in the first mentioned cluster (pct.1) and in the second mentioned cluster (pct.2), and the FDR adjusted *P* value (p_val_adj). Associated with Figure 1—figure supplement 1, Figure 3, Figure 4, Figure 4 - figure supplement 1.

Supplementary file 2

Gene ontology (GO) analysis of DEGs. (A) GO analysis of genes upregulated in Prolif SC *versus* SC1+SC2 in neonatal AP. (B) GO analysis of genes upregulated in Prolif *Pou1f1*^+^ cells *versus* Somato in neonatal AP. (C) GO analysis of genes upregulated in Prolif Cortico *versus* Cortico in neonatal AP. (D) GO analysis of genes upregulated in Gonado Prog *versus* SC1+SC2+Prolif SC in neonatal AP. (E) GO analysis of genes upregulated in neonatal *versus* adult SC1+SC2+Prolif SC. (F) GO analysis of genes upregulated in neonatal damaged *versus* control Prolif *Pou1f1*^+^ cells + *Pou1f1* Lin Prog. (G) GO analysis of genes upregulated in neonatal control *versus* damaged SC1+SC2+Prolif SC. Analyses were performed using Reactome overrepresentation analysis for (A-C) and (E-G). Pathway name, number of mapped identifiers that match the pathway (#Entities found), total number of identifiers in the pathway (#Entities total), total entities in the pathway/total number of entities for the entire species (Entities ratio), *P* value, FDR and -log10(FDR) are shown in the columns. Analysis was performed using Gorilla for (D). Pathway name, *P* value, FDR q-value, enrichment score [(b/n)/(B/N)] and -log10(FDRq) are shown in the columns. N, total number of genes; B, total number of genes associated with a specific GO term; n, number of genes in the top of the user’s input list or in the target set when appropriate; b, number of genes in the intersection. Associated with Figure 1—figure supplement 1, Figure 3, Figure 4, Figure 4 - figure supplement 1.

Supplementary file 3

Overview of medium components in PitOM and RSpECT, used for organoid culturing.

Supplementary file 4

Overview of primary and secondary antibodies, used for immunofluorescence staining.

Supplementary file 5

Overview of primers, used for RT-qPCR.

